# Shortfalls and opportunities in terrestrial vertebrate species discovery

**DOI:** 10.1101/2020.10.23.352690

**Authors:** Mario R. Moura, Walter Jetz

## Abstract

Meter-resolution imagery of our world and myriad biodiversity records collected through citizen scientists and automated sensors belie the fact that much of the planet’s biodiversity remains undiscovered. Conservative estimates suggest only 13 to 18% of all living species may be known at this point ^1–4^, although this number could be as low as 1.5% ^5^. This biodiversity shortfall ^6,7^ strongly impedes the sustainable management of our planet’s resources, as the potential ecological and economic relevance of undiscovered species remains unrecognized ^8^. Here we use model-based predictions of terrestrial vertebrate species discovery to estimate future taxonomic and geographic discovery opportunities. Our model identifies distinct taxonomic and geographic unevenness in future discovery potential, with greatest opportunities for amphibians and reptiles and for Neotropical and IndoMalayan forests. Brazil, Indonesia, Madagascar, and Colombia emerge as holding greatest discovery opportunities, with a quarter of future species descriptions expected there. These findings highlight the significance of international support for taxonomic initiatives and the potential of quantitative models to aid the discovery of species before their functions are lost in ignorance ^8^. As nations draw up new policy goals under the post-2020 global biodiversity framework, a better understanding of the magnitude and geography of this known unknown is critical to inform goals and priorities ^9^ and to minimize future discoveries lost to extinction^10^.

## Main Text

Previous studies have tackled this challenge through a range of extrapolation techniques using species discovery curves and expert opinion ^1–3,11^, but with limited detail beyond global/continental taxon percentages or counts ^12,13^. Here we use the effects organismal characteristics have on discovery probability to provide taxonomic and geographic specificity of future species discovery ^14–17^. For example, take one of the largest extant birds, the emu *Dromaius novaehollandiae*, which was described in 1790 in a time and region with limited taxonomic activity; centuries before a small, elusive frog species *Brachycephalus guarani*, discovered in 2012 in Brazil (Fig 1a). The difference between the two species matches previous insights about the effects of body size, range size and taxonomic activity on discovery ^16,18–20^. We extend this comparison to eleven biological, environmental, and sociological attributes in a ‘time-to-event’ model framework to estimate the probability a given species’ discovery (event) over time ^21,22^ (Supplementary Information).

**Fig. 1.**
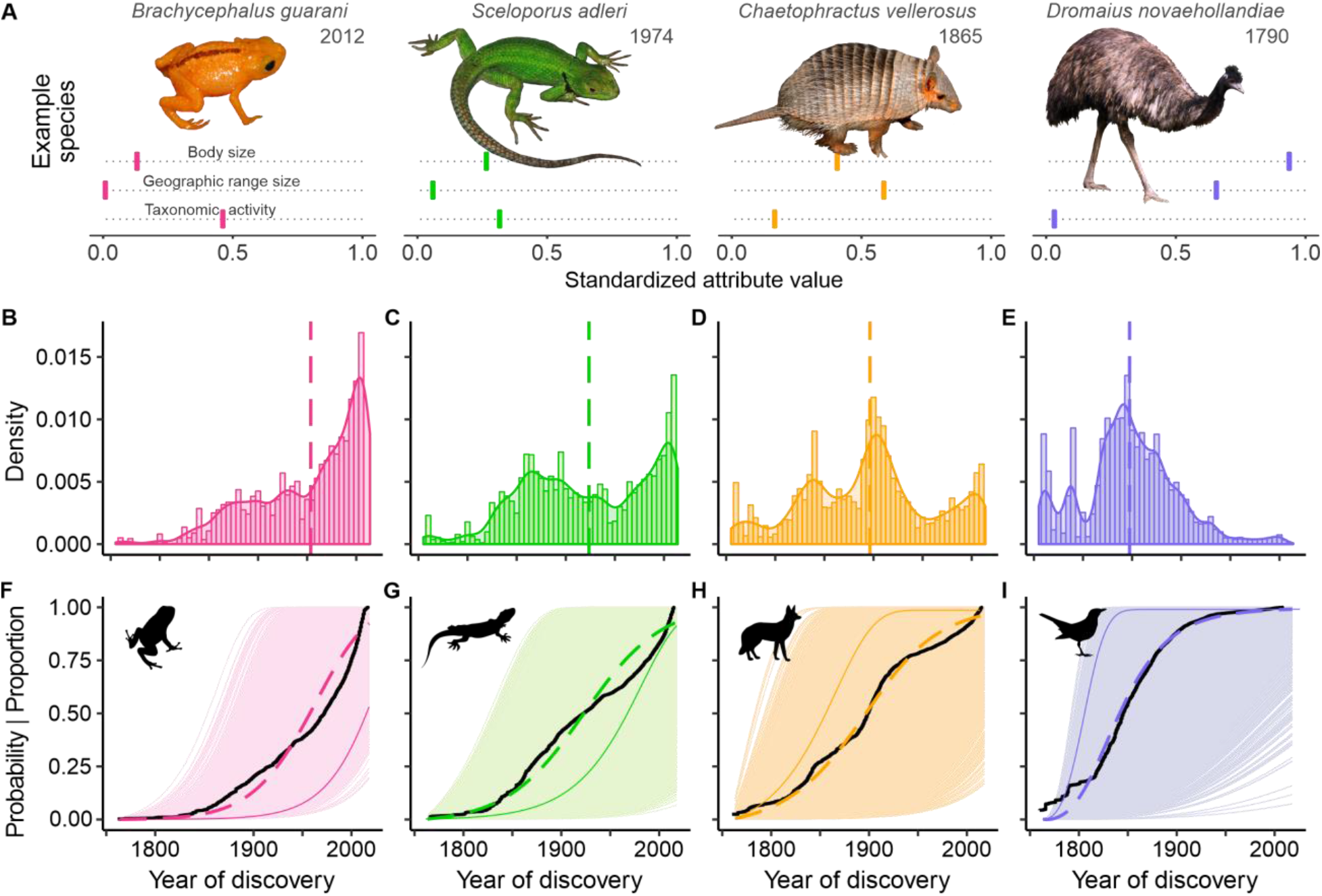
Variation in observed and predicted discovery trends for the years 1759-2014 across the four terrestrial vertebrate groups. (A) Example species and their attributes (standardized to vary from 0 and 1 in each group, separately). (B-E) Variation in species descriptions over time. Vertical dashed lines indicate the year in which 50% of the known species were described. (F-I) Time-to-event model-based predictions of discovery probability for each species (light colours; solid coloured line representing example in A), and average trends across all (coloured dashed-line). Black lines show the empirical cumulative growth of described species described across time (expressed as proportion of known species).

For all extant species, the modelling framework provides predicted discovery year and discovery probability at the present (herein 2015) based on the weighted importance of each assessed attribute. In our earlier example, *B. guarani* (described in 2012) had a 49% (95%CI: 37-64%) chance of discovery by 2015 given its attributes. Conversely, the discovery probability for the emu exceeded 50% already in 1759, increasing to 100% by 2015 (95%CI: 100-100%). When applied to 32,172 species of amphibians, reptiles, birds, and mammals, the predicted discovery curves match observed differences in temporal description patterns in the four taxa. Most bird species saw high discovery probabilities early on, matching the median avian description year of 1845. In contrast, half of all amphibian descriptions occurred after 1972, and modelled discovery curves accordingly show a slow increase (Fig. 1).

Species’ body size, geographic range size, and taxonomic activity strongly affect variation in discovery probability, but terrain and environmental conditions also matter ^16,19,23^. Species tend to have higher discovery probability if they are large-bodied, wide-ranged, located in cold climates or characterized by, at the time, low taxonomic activity or low human density (Fig. 2). The magnitude, and sometimes also direction, of effects differs somewhat among the four groups. It also varies across time, reflecting developments in taxonomists’ modes and toolbox, as well as changes in the kinds of species left to be discovered (Fig S3–S6). For example, among more recently discovered species, body size or human density have lost predictive strength. In amphibians, higher elevations are less of a constraint on discovery probability than in the past, whereas the recency of mammal discovery continues to be associated with higher elevations. Among bird species described since the mid-20^th^ century, wetter locations have yielded later discoveries, but not so prior to that time. Notably, in amphibians and reptiles, clades and regions with more active taxonomists remained those with greatest discovery potential. This highlights how gaps in taxonomic expertise continue to limit our recognition of species.

**Fig. 2.**
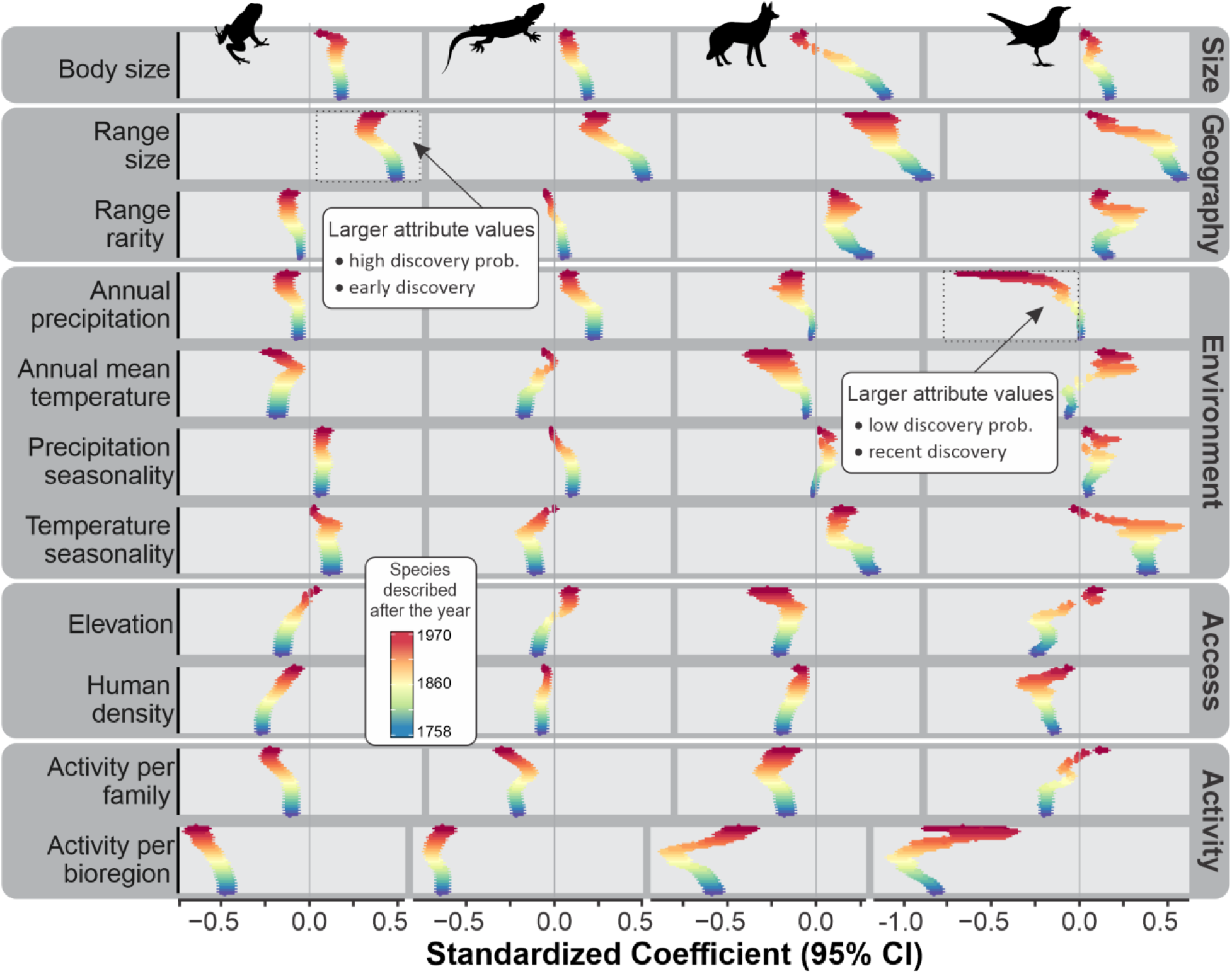
Joint effects of species-level attributes on discovery probability over different time periods. Standardized coefficients above 0 indicate that species with high values for a given attribute had higher discovery probability (prob.) and thus were likely discovered early on. Negative standardized coefficients mean high attribute values depressed discovery probability and delayed discovery. The vertical colour gradients illustrate the variation in coefficient from all (bottom) to more recently described species as species are successively removed from the analysis. Coefficients include 95% confidence intervals as horizontal bars.

Averaged across species in an assemblage or clade, the divergence of modelled discovery probability from 100% informs the portion of species yet to be discovered given past modes of description. Among vertebrate clades with >5 species, South American shrew opossums (Paucituberculata), dibamids, geckos and relatives, wall lizards and other lacertids emerge as having the greatest relative undescribed diversity (Fig. 3). Scaled by groups’ species count, the models identify several frog clades, geckos and iguanas and their relatives, and snakes as the vertebrate groups with the highest expected number of future species discoveries. Among mammals, rodents and bats feature in the top ten higher-level taxa, partly reflective of the already large described diversity of these groups ^24^. Cross-validation with holdout data on past discoveries indicate a strong predictive power of our framework and suggest robustness across groups (Extended Data, Figs S8, S14). While some of the identified taxonomic discovery hotspots are not unexpected, our evaluation across all terrestrial vertebrates offers a transparent quantitative comparison in support of taxonomic research priorities.

**Fig. 3.**
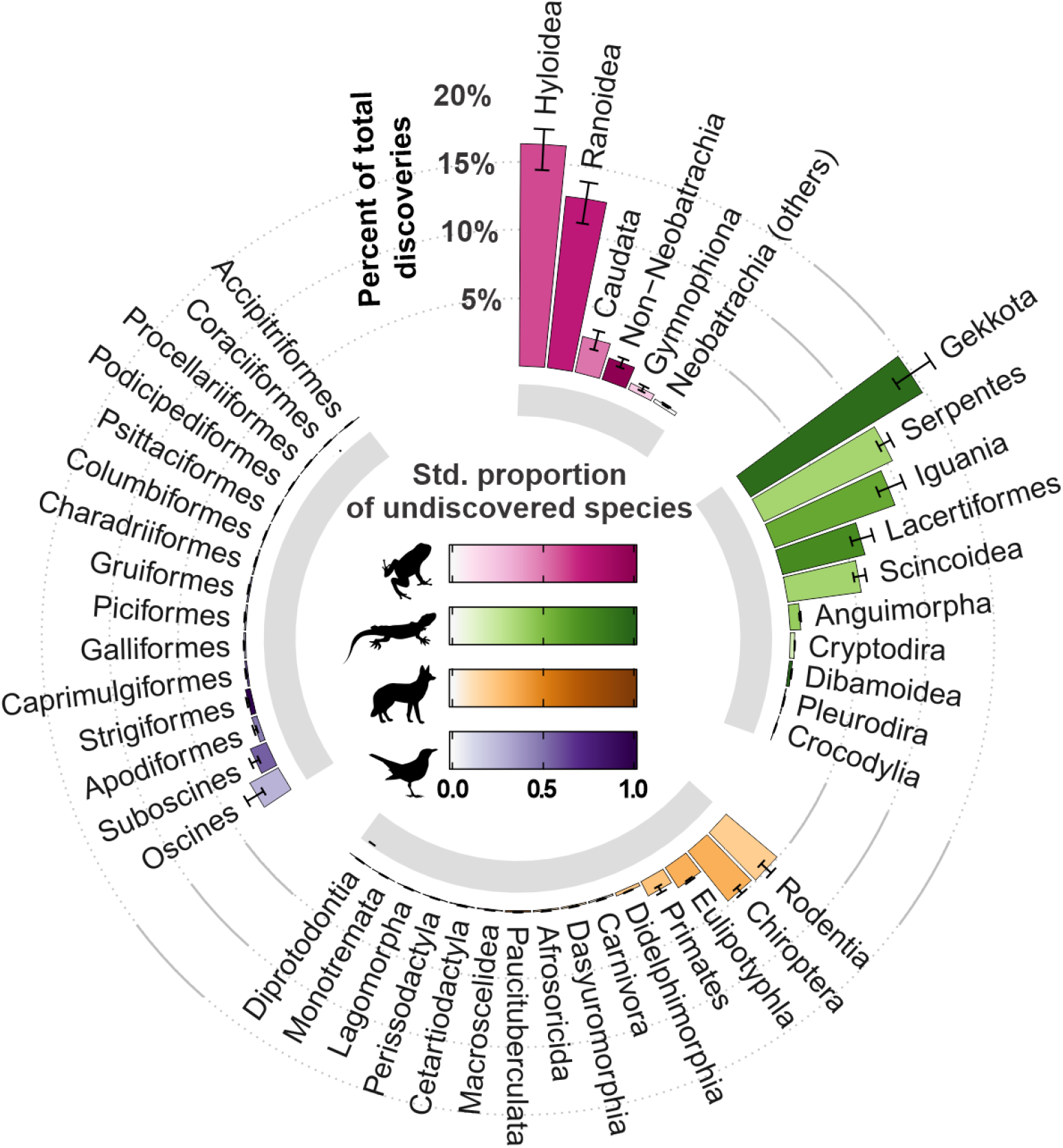
Predicted future discovery potential across major terrestrial vertebrate taxa. Bar height indicates the percentage of all future terrestrial vertebrate discoveries predicted to occur in the taxon, with error bars indicating 95% confidence intervals. Bar colours show the proportion of undiscovered species within each vertebrate class, standardized to vary from 0 to 1 (see Supplementary Information). See Figs S16–S19 (Extended Data) for the full set of discovery metrics at family level.

Both the facilitators of species discovery – such as fieldwork and systematics initiatives – and the drivers of undiscovered species’ demise – such climate- and land-use change – are strongly place-based ^25,26^. We therefore extended our discovery predictions to geographic space. We mapped attribute-driven discovery probabilities across species distributions while applying a subsampling procedure accounting for range-size driven variation in representation ^27^. As the locations with highest number of expected future discoveries (Fig. 4), we identify the Tropical Andes and the Atlantic Forest, the Eastern Afromontane and West African Guinean Forests, Madagascar, and the Western Ghats, Sri Lanka, Indo-Burma, and Philippines and New Guinea Forests. Projected unknown species richness covaries with extant species richness (Spearman *r* = 0.87–0.90), but discovery hotspots also included locations with relative limited extant diversity such as the Southern Andes and Caatinga region. Validations with observed discoveries strongly support these projected spatial discovery patterns across different spatial resolutions (Spearman *r* = 0.71–0.92 for all groups, see Extended Data Figs S9–S12, S15).

**Fig. 4.**
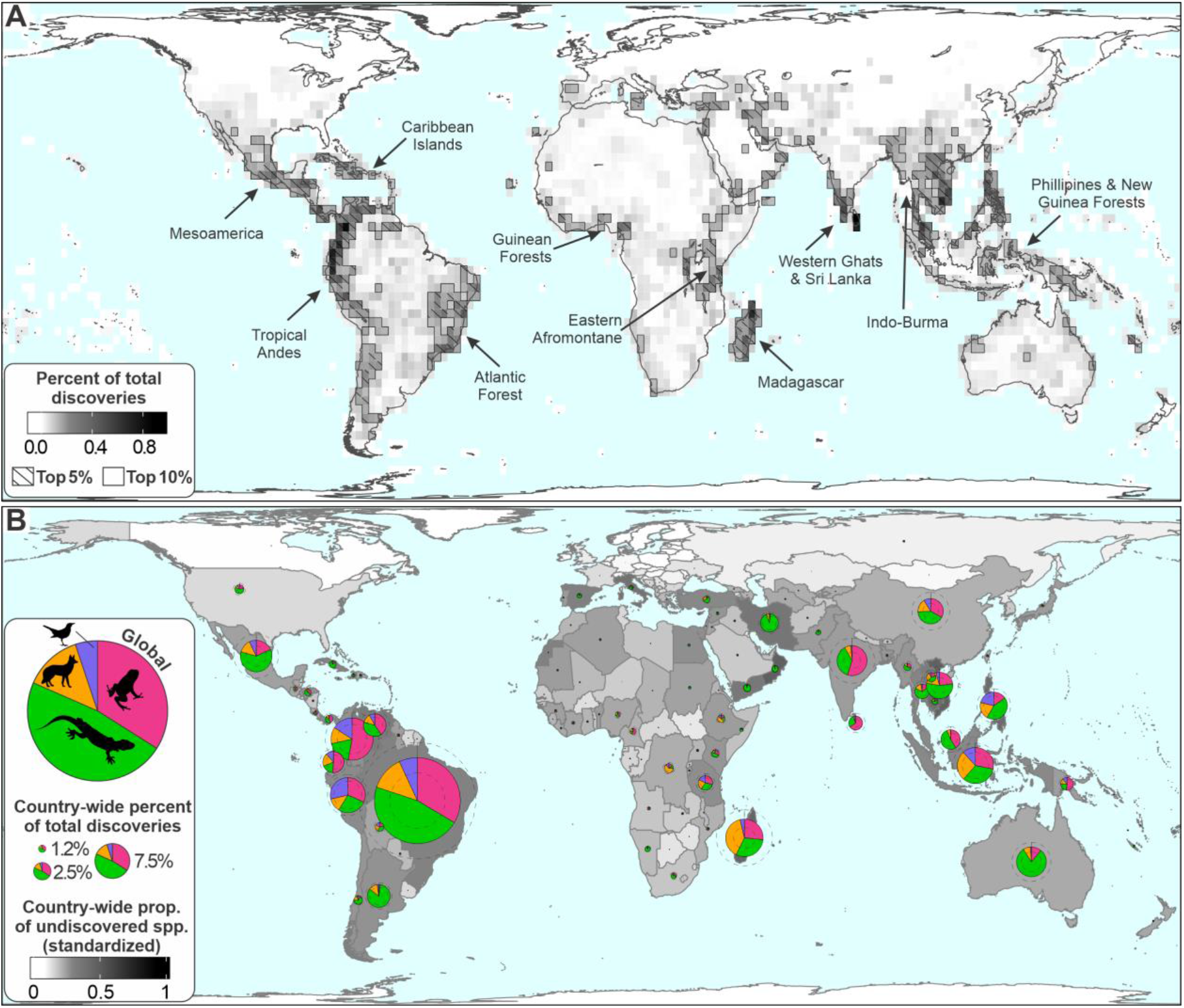
Global variation in predicted discovery potential, quantified as the percent of all global terrestrial vertebrate discoveries predicted to occur in a region. (A) Variation across 220km grid cells, standardized to percent of total discoveries. (B) Variation among countries, with colours showing mean discovery potential, expressed as country-wide proportion (prop.) of undiscovered species (spp.) and standardized to vary from zero to one. Pie charts illustrate the predicted distribution of discoveries among the four vertebrate classes in each country (“Global” in legend shows the global pattern); pie chart size indicates the country-wide total, with dashed grey lines indicating the 95% confidence interval. See Figs S20–S24 (Extended Data) for maps of the discovery metrics at different spatial resolutions.

Expertise, support, and incentives for future discovery are ultimately tied to nations – the stewards of these unknown biological resources. Aggregating our spatial estimates to countries highlights several South American and South Asian nations and Madagascar as countries with highest projected future discoveries, i.e. greatest “discovery debt” or conversely, “biodiversity reward” (Fig. 4, and Fig. S25, Extended Data). These countries are home to a large diversity of taxa with attributes indicative of low discovery rate to date, and thus likely contain many expected future discoveries. Brazil stands out with multiple diversity centres across its large area, holding ca. 10.5–10.8% of all projected future discoveries, which largely coincides with the 10.8–13.4% of ant genera discovery projected for this country ^13^. Other top discovery debt countries for terrestrial vertebrates include Indonesia (5.0–5.7%), Madagascar (3.9–4.9%), and Colombia (4.1–4.4%).

Reflecting their class-wide high discovery potential, reptiles constitute the greatest portion of this future prospect. Undiscovered reptiles are expected in more arid regions, such as Australia, Iran, and Argentina, correlating with existing centres of reptile diversity and endemism ^28^. In the tropics, many countries owe most future discoveries to amphibians, particularly in southern Asia and northern South America (Extended Data, Fig. S25). Discovery potential for mammals is more limited and concentrated in recent description hotspots such as Madagascar. Compared to other taxa and reflecting their flattened description curve (Fig. 1), the discovery potential for birds is low, but given past trends the model estimates further discoveries especially in Peru, Colombia, Brazil, and Philippines.

Research on quantifying species discovery shortfalls is by definition imprecise, with absolute estimates of undiscovered species differing by orders of magnitude ^1,3,5^. We focused on the geographic and taxonomic differentiation in discovery potential and used a well-known subset of global biodiversity to develop a generalizable framework to address this challenge. We do not expect that our discovery projections will hold up in exact form. The present estimates are a direct reflection of past description processes and their correlates, and any forward interpretation therefore needs to recognize intrinsic limitations. Despite ongoing calls for taxonomic standardization and stability ^29,30^, species also represent scientific hypotheses that are sometimes revisited, refuted, or revalidated ^31,32^. Our models therefore are not able to distinguish operational definitions of valid species and the potential heterogeneous associations arising from variable practices around, e.g., recognizing cryptic species or splits ^33^. There may also be parts of the multivariate predictor space that lack data to inform the model and thus miss actual discovery opportunities. Nevertheless, extensive model validations confirmed a strong predictive ability for species discoveries and highlight the potential to increase discovery rates through the use of quantitative frameworks such as the one presented.

Our findings indicate that discovery gaps hinder the safeguarding and realization of biodiversity for certain kinds of species and for select places and countries much more than others. We show that specific countries require increased capacity and support to address this challenge. After centuries of efforts by biodiversity explorers and taxonomists, the catalogue of life still has too many blank pages. Extending the presented approach to other taxa has the potential to underpin taxonomic research initiatives that help speed up discovery before species are lost in ignorance ^34^. With discussions of the Post-2020 Global Biodiversity Framework ongoing, we urge intergovernmental recognition of the unevenness in this knowledge shortfall and of the growing scientific opportunities to address it.

## METHODS

We used species-level biological, environmental, and sociological attributes to parameterize time-to-event models of discovery probability across time in birds, mammals, amphibians, and reptiles ^21^. The models provided for each species an estimated probability of discovery in the present time given its attributes. We used these species predictions to characterize the major taxonomic and geographic groups they are part of for their future discovery potential. Specifically, for major taxa and assemblages worldwide the central tendency (geometric mean) of model-driven discovery probability of its member species informed an estimate of the proportion of their known vs. yet to discovered species richness.

### Species data

We compiled trait data for nearly all extant species of terrestrial vertebrates ^28,35^, excluding those with uncertain geographic distribution, that is, species with occurrence reported only at the level of administrative units – country, states, etc – without precise location. We also excluded species described after 2014 to minimize potential biases from their potentially incomplete geographic characterization. Overall, our dataset comprised 32,878 species of terrestrial vertebrates: 7202 species of amphibians, 10,004 of reptiles, 5679 of mammals, and 9993 of birds (Data S1). Taxonomic nomenclature followed the same adopted in original data sources ^28,36–39^.

### Species-level attributes

Previous studies have shown that recently described species often have smaller body sizes and narrower geographic range compared to those described earlier ^16,17^. But species detectability and discovery may be affected by a range of other attributes, such as the environmental and socioeconomic conditions where the species occurs, and the taxonomic knowledge of a given taxon or region ^16,17,40^. We considered a total of eleven putative correlates of discovery probability (see Data S5):

i. Body size. Larger animals are easier to detect thus being described first ^16,17,23,41^. Information on maximum body size was compiled from available data sources: amphibians ^42^, reptiles ^43–47^, mammals ^38,48^, and birds ^49^. We complemented these data sources by inspecting the literature for body size information of species without data. In the end, we obtained body size data for 6869 (95.2%) species of amphibians, 9852 (99.7%) of reptiles, 5208 (91.4%) of mammals, and 9123 (91.3%) of birds. For amphibians, body size was represented by snout-vent length (in mm, anurans) or total length (in mm, caecilians and salamanders). Reptiles had their body length measures converted into masses (g) using taxon-specific allometric equations available in the literature for squamates ^43^ or here developed for chelonians (Supplementary Information, Table S1). Birds and mammals had their body size represented by masses (g), as provided in the available data sources ^38,49^. For 870 bird species with missing body size data, we used the genus-level mean body size as originally provided in ^49^. For the remaining species without body size data, we performed a phylogenetic imputation using the fully-sampled global phylogenies made recently available ^37,38,50^ in concert with the R package *Rphylopars* ^51^. We discarded seven reptile species due to the lack of both body size and phylogenetic data (*Agama congica*, *Bothrochilus montanus*, *Gerrhosaurus intermedius*, *Hemidactylus benguellensis*, *Leptotyphlops lepezi*, *Rhoptropus benguellensis*, *R. montanus*). To account for uncertainty in the fully sampled phylogenies, we used 100 trees of each vertebrate group to perform imputations. We loaded the named species-level data (including missing values) and ran *phylopars* function using Brownian motion (BM) as model of trait evolution. We then matched the imputed values back to the taxonomy and summarized the BM imputations across the 100 trees as medians for each species. All body size measures were log_10_ transformed before phylogenetic imputations. Overall, we imputed body size for 333 species of amphibians, 23 of reptiles, 492 of mammals. We assumed intraspecific variation in body size to be negligible relative to interspecific variation.
ii. Geographic range size. Widely distributed species tend to be locally abundant ^52,53^ and are therefore easier to find, being described earlier than narrowly distributed species ^16,17,23,41^. We overlaid expert-based extent-of-occurrence range maps of each species with an equal-area grid of 110 ×110 km cell size. Range maps were extracted from ^39,54^ for amphibians, ^28^ for reptiles, ^35,38,54^ for mammals, and ^35,36^ for birds. Range size was then measured as the number of grid cells intersected by each species. Only the native and breeding range of species were considered for these computations. Presence of a species in a grid cell was recorded if any part of the species distribution polygon overlapped with the grid cell.
iii. Range rarity. Biodiversity researchers may prefer to work in areas with many or geographically rare species, and describe first the species from those areas ^25,55^. A commonly used metric of rarity is the total range size rarity, also called endemism richness, defined as the sum of the inverse range sizes of all species present in a place ^56^. To represent rarity at the species-level, we used the average endemism-richness within each species’ range. However, grid cells (regions) that currently harbour many and/or rare species had not always been known as such, since known richness and endemism patterns may change through time as species descriptions progressed. To better capture the variation in range rarity across time, we computed the endemism richness using only species described from 1758 to *x*, where *x* varied from 1758 to 2014. Then, for each species, we computed the average known within-range endemism richness at the year it was described.
iv. Annual precipitation. Early descriptions dates are on average low for species occurring in Europe, North America, and western Asia ^57^. These later regions have received substantial taxonomic effort, which explains their higher levels of inventory completeness relative to tropical and desert-like environments ^25^. Thus, it is reasonable to consider that early naturalists were trained in temperate regions and therefore they explored first species from relatively dry regions ^16,58–60^. We calculated the average annual precipitation in each equal-area grid cell at 110 ×110 km of spatial resolution and then computed the average within-range annual precipitation for each species using the 1 km climatic layer from ^61^. Computations were performed in the R software ^62^ using the *extract* function of ‘raster’ package ^63^.
v. Annual mean temperature: Following the reasoning aforementioned, early naturalists were trained in temperate regions and therefore they explored first species from cold regions^16,58,59^. We calculated the average temperature (annual mean) in each equal-area grid cell at 110 ×110 km of spatial resolution and then computed the average within-range temperature for each species using the 1 km climatic layer from ^61^.
vi. Precipitation seasonality. Early naturalists were trained in temperate regions and therefore they explored first species from high seasonal regions ^23,58^. We calculated average within-range precipitation seasonality (coefficient of variation) for each species using the 1 km climatic layer from ^61^.
vii. Temperature seasonality. Early naturalists were trained in temperate regions and therefore they explored first species from high seasonal regions ^16,58,59^. We measured average within-range annual temperature seasonality (standard deviation) for each species using the 1 km climatic layer from ^61^. Although climatic conditions might not have been necessarily stable over the last centuries, we assumed that average conditions from 1979-2015 represented similar climatic conditions from the 18th to 20th centuries. We argue that averaging climatic variables within species geographic range may both dilute the temporal variations in local climate and avoid the uncertainty associated with extrapolating fine resolution climatic data to past times of low density (or even absence) of weather stations.
viii. Elevation. Mountainous regions might have limited accessibility, likely impeding early species descriptions from higher elevations ^25,40^. Although early taxonomists and naturalists likely explored low elevation regions first, we avoided computations of minimum within-range elevation since species with coastal distribution could show biased values that do not necessarily reflect the most common elevation where they occur. We computed the mean elevation within each equal-area grid cell at 110 ×110 km of spatial resolution, using the 1 km global topography layer from ^64^. We then extracted the average within-range elevation for each species.
ix. Human density: A species may be described if human population density within its geographic range surpass a detectability threshold that enhance its discovery probability^23,40^. Such detectability is expected to be low before the species description (too few or even no humans overlapping the species geographic range) and irrelevant after its discovery (human density might change but the formal description already happened). Thus, an informative measure of geographic range overlap with human settlements should consider the year of species description. Earlier descriptions occurred at times of low human density and much likely involved easily detectable species, whereas more recent descriptions have coincided with times of high human density. We therefore expect a positive association between human density and year of description. To quantify the influence of humans on species description, we computed the average human density within the species’ range at the exact year of its description (for all species described after 2000), or at the closest decade (for all species described before 2000). Historical data on human density data was obtained from ^65^ at the spatial resolution of 5 arc-min (~10 km).
x. Activity per family. In general, taxonomists tend to discover the ‘obvious’ first and ‘obscure’ later. Species like the emu (Fig. 1) were quickly noticed by taxonomists, even in times of low taxonomic activity (relative to current trends). However, other species exhibit high conspicuousness and likely required more attention to be taxonomically noticed. This level of attention is what we refer here as taxonomic activity. In increasing the number of active taxonomists per family, a potentially inconspicuous or cryptic species may receive enough attention to have its discovery unveiled by taxonomists. Otherwise, a species may remain unknown while the taxonomic activity within its family is kept low. Similarly to human population, the number of taxonomists has increased over time, with fewer taxonomists authoring early species descriptions ^66^. Therefore, we expect species described long ago to show low within-family taxonomic activity, and consequently high discovery probability. We used the number of authors of each species description as a proxy for taxonomic activity. We standardized the surnames included in the authority name of each species to lowercase letters without special characters. For each species, we identified the taxa described in the same family and year as the focal taxon and then calculated the aggregate number of unique active taxonomists. Because the number of active taxonomists in a given year is expected to increase as more species are described in that year, we standardized this measure by dividing it by the number of within-family descriptions in the respective year. Computations were performed using the *stringr* ^67^ and *splitstackshape* ^68^ R-packages.
xi. Activity per bioregion. In a similar way to the influence of taxonomic activity per family, a new species may go unnoticed while the number of taxonomists working within its geographic range remain too low or even non-existent. Under low levels of taxonomic activity, those easy to find species are likely described first. In other words, species with high ‘taxonomic conspicuousness’ may require eyes from more taxonomists. Following the trend of increasing the number of taxonomists per species over time ^66^, we expect early described species to show low within range taxonomic activity, in contrast to recent described species. These computations followed those for the taxon level, but with species instead subset by geography. Specifically, we used the biogeographical realm and biome classification proposed in ^69^ to compute the percentage overlap between species geographic ranges and realm-biome combination, or bioregion ^70^ (e.g. Tropical and subtropical moist broadleaf forests in the Neotropics). Each species was classified as typical of a given bioregion if it either occurred in at least 25% of the bioregion or the bioregion intersected with at least 25% of its geographic range (species could be typical of multiple bioregions). For each species, we then selected its ‘typical’ bioregion and extracted all other species described in the same year and co-occurring in the same bioregion as focal taxon. We then summed the number of unique taxonomist names that described species within the selected bioregion and divided it by the number of within-bioregion descriptions to obtain a measure independent of the number species descriptions. We recognize that our metrics of taxonomic activity might be affected by duplicated names (same taxonomist entered with the different surname) or homonyms (taxonomists with the same surname), but given the spatial, taxonomic, and temporal constraints applied, we expect this issue to be negligible.

Since all our predictor variables vary over many orders of magnitude, we log_10_ transformed them to reduce their skewness. We examined multicollinearity of the predictor variables using the Variance Inflation Factor (VIF). Predictors holding VIF values > 10 are regarded as having high multicollinearity and should be excluded from the model ^71^. As none of our predictors achieved VIF > 5 (Supplementary Information, Table S2), we kept all of them for the subsequent analysis. VIF computations were performed with the ‘usdm’ R package ^72^.

### Time-to-event models

We used time-to-event analysis, also known as survival analysis ^21^, to assess the effect of species-level attributes on the description rates observed in a given vertebrate class. Time-to-event analysis is commonly used in the medical, engineering, and social sciences to assess factors influencing the probability of an event (e.g., death, mechanical failure, getting a job), but has also been used in ecological studies ^22,73,74^. In our analysis, the event of interest is the species description date and the measure of time the number of years passed until this date since 1758, the beginning of modern taxonomy through Linnaeus ^75^. Although our species data covers the period from 1758 to 2014, we did not use species described in the year 1758 itself to avoid the large number of descriptions due to Linnaeus work ^75^. Overall, our class-level time-to-event models were informed by 7,185 species of amphibians, 9,889 of reptiles, 9,557 of birds, and 5,541 of terrestrial mammals described between 1759 and 2014.

Specifically, we modelled time-to-description using Accelerated Failure Time (AFT) model, which is a parametric time-to-event model to evaluate covariate effects on the acceleration/deceleration of the probability of an event ^76^. The output of the AFT model includes the probability of a given species to have remained unknown across time, i.e. its survival probability. We define the 1 minus species survival probability as species discovery probability (Fig. 1). This discovery probability is always increasing; as time moves forward, we accumulate chances to discover an unknown species.

We initially ran a model selection procedure to identify the family error distribution that is best suited to our time-to-description variable ^77^. For each of the four vertebrate classes, we built null AFT models (Time to event ~ 1) using six different family error distributions (Exponential, Weibull, Log-normal, Log-logistic, Gamma, and Gompertz) available in the *flexsurvreg* function from the ‘flexsurv*’* R package ^78^. We then identified the model offering the best error family distribution using the Bayesian Information Criterion (BIC) ^79^. Once the best family error distribution was selected, we proceeded with the subsequent analysis using the predictor variables.

Given the high number of possible models using all predictor combinations (2^11^ – 1 = 2047 models), it may be difficult to find an overwhelmingly supported model because the best predictors (if any) will have their importance diluted among multiple models ^80^. Hence, to incorporate the uncertainty around the variable selection procedure into our model coefficients, we passed the predictors through a model averaging procedure ^81^ and for each possible AFT model obtained the standardized coefficients. Computations performed using the ‘MuMIn’ ^82^ and ‘stats’ ^62^ R packages.

### Species-level predictions

For every possible AFT model we computed the discovery probability of each species in year 2015 (herein considered as ‘present time’, Data S1). To relate this value to observed description dates, we used a threshold of 0.5 to convert the estimated discovery probability for a given time step into a predicted description and extracted the corresponding year as predicted description date. We note that the discovery curves returned by an AFT model have the same shape but a different position along the time axis, the latter being determined by the covariates of each species. Because of that, using a different threshold for the binary conversion does not affect the slope of the relationship between the observed and estimated description dates, only the intercept changes if a different threshold value is applied. This procedure yielded for every species and each of the 2,047 AFT models i) a discovery probability in year 2015 and ii) a predicted description date. We weighted these metrics by the relative BIC weights (wBIC) of their models to arrive at species-level discovery metrics used in subsequent analyses. Computations performed using the ‘MuMIn’ R package ^82^. All species-level estimates are available through Data S1 file.

### Taxon-level predictions

We used the species-level predictions of discovery probabilities to characterize individual families and higher-level groupings for their potential for future discoveries. Specifically, we estimated the taxon-level proportion of known species to date (*PropKnown*) as the central tendency (geometric mean) of discovery probability of its member species ^22^. We also obtained the known species richness (*KnownSR*) per taxon, and used both *PropKnown* and *KnownSR* to calculate total richness (*TotalSR*) through a rule of three: *TotalSR* = *KnownSR* × 100 / *PropKnown*. If the *PropKnown* of a given taxon was 100%, then all species were expected to be described in the respective taxon, and *TotalSR* = *KnownSR*. If *PropKnown* was < 100%, then *TotalSR* > *KnownSR*, and the difference between these variables represented the unknown species richness: *UnknownSR* = *TotalSR* – *KnownSR*. The proportion of unknown species (*PropUnknown*) was given by 1 minus *PropKnown*. We highlight that *PropKnown* (our measure of central tendency), represents a snapshot of the current biodiversity knowledge for a given species sample. In approaching saturation of species discovery in a sample, *PropKnown* is expected to shift towards 100%. We only computed these metrics for families and higher-level groupings holding five or more species.

For mammals and birds, we used taxonomic orders as the highest-level taxonomic rank, although we split Passeriformes birds into Oscines and Suboscines. In amphibians and reptiles, orders are highly uneven in size, which led us to use a more informative higher-level grouping for them. For amphibians, we kept the orders Gymnophiona (caecilians) and Caudata (salamanders and relatives), and followed ^83^ to divide the order Anura into four groups: (i) non-Neobatrachia (some primitive anuran families), (ii) Hyloidea taxon within Neobatrachia, including most frog species from the Neartic and Neotropic realms, (iii) Ranoidea taxon within Neobatrachia, including most frog species from the Afrotropical, Paleartic, Indo-Malay, and Australasia realms, and (iv) other Neobatrachia (a non-monophyletic set of Neobatrachian families not included in Ranoidea or Hyloidea). For reptiles, we kept the order Crocodylia (alligators and relatives), divided the order Testudines in the suborders Pleurodira (side-necked turtles) and Cryptodira (hidden-necked turtles), and followed ^84^ to split the order Squamata into seven groups: Gekkota (geckos and relatives), Iguania (iguanas, chameleons, and relatives), Scincoidea (skinks and relatives), Lacertoidea (teiids, lacertids, amphisbaenians, and relatives), Anguimorpha (glass lizards, monitors, and relatives), Dibamidae (dibamids or blind skinks, also referred as Dibamoidea), and Serpentes (snakes).

The model validation (see below) indicated that our framework satisfactorily identified the relative potential of taxa to hold unknown species, but it underestimated absolute values of *UnknownSR* and *PropUnknown* per taxon (Supplementary Information, Table S4; Extended Data, Fig. S8). Thus, we standardized both measures of discovery potential. For each vertebrate class, we divided the *UnknownSR* by the total number of estimated discoveries (i.e., sum of *UnknownSR* across taxa) to provide the estimated percent of total discoveries. The proportion of unknown species (*PropUnknown*) per taxa was standardized to vary between 0 and 1, by first subtracting the minimum observed for each vertebrate class and then dividing by the respective range of *PropUnknown*. The value of 1 indicated the taxon with the highest proportion of unknown species (whatever such number might be), and not necessarily a taxon with 100% of unknown species. All taxon-level estimates are available through Data S2 files.

### Assemblage-level predictions

We followed the same rationale we used to for taxon above to estimate the variation in future discovery potential in geographic space. Specifically, we considered the proportion of species that remain to be discovered in an assemblage, *PropUnknown*, an emergent property of its species members and their attributes. This approach follows the growing recognition in trait biogeography and macroecology of the species-level drivers of larger-scale patterns ^14,22,85–87^. We used the equal-area grid cell species distribution data (see above) to derive species lists for each the four vertebrate classes for assemblages of 220, 440, and 880 km grid cell size, discarding all assemblages with less than five species. For each assemblage we then calculated *PropUnknown* and *UnknownSR* based on the discovery probabilities of its member species, the same way we did at the taxon-level.

Different to the by-taxon characterization, however, in the assemblage patterns wide-ranging species are overrepresented, since those are counted multiple times throughout grid cells ^36,88^. Considering that widely distributed species tend to be described first ^16,17,41^, this can bias assemblage measures towards lower *PropUnknown*. This unevenness in the representation of wide-ranging species can be controlled through a random subsampling approach that provides range-size controlled estimates of aggregate measures at the assemblage level ^27^.

Briefly, the subsampling algorithm we applied considers the random extraction of *x* grid cells belonging to a given species’ range. If the species’ range was smaller than *x*, then, all grid cells are extracted for that species. The *x* here is analogous to the pseudoreplication level of a dataset. If *x* equals 1, then the geographic range of all species will be subsampled to show only one grid cell per species. The subsampling algorithm was applied to all species in a single iteration, and the subsampled geographic ranges were then overlapped for the extraction of the *PropUnknown* and *UnknownSR*. These computations were performed 100 times, and the mean value across iterations was extracted for each grid cell to represent the geographic pattern of the respective aggregate measure under the *x* level of pseudoreplication. Additional details on this subsampling algorithm are available in ^27^. In this study, we used seven different levels of *x*: 1, 5, 10, 50, 100, 200, and 500 grid cell occurrences per species, also including the observed pattern using all grid cell occurrences per species. Computations performed using ‘data.table’ R package ^89^.

As in many ecogeographical investigations ^90^, this study focuses in the variation of species-level attributes across space to describe and explain biodiversity patterns. Given the dominant unavailability of spatially varying data on species-level attributes, most ecogeographical studies – including this one – assume species-level attributes to be spatially constant. While the incorporation of spatially varying covariates through hierarchical modelling remains an open avenue in trait biogeography^91^, we argue that it has limited influence in our results for two major reasons. First, most species in our dataset are not widely distributed, which implies less potential for biological attributes to vary in space. For instance, 50%, 70%, and 88% of species in our dataset occupy ≤4 grid cells at respectively 220, 440, and 880 km of spatial resolution (See Data Availability for raw data). Second, the subsampling algorithm we applied reduces the overrepresentation of widely distributed species, whom are the ones with highest potential to show spatially varying biological attributes.

We standardized *UnknownSR* and *PropUnknown* per assemblage (Supplementary Information, Table S4; Extended Data, Fig. S8) in the same way as done for taxa. We divided the *UnknownSR* per assemblage by the sum of *UnknownSR* across assemblages to get the estimated percent of total discoveries, and standardized *PropUnknown* to vary between 0 and 1, with 1 indicating the assemblage with the highest proportion of unknown species (whatever such number might be). We note that adding up the *UnknownSR* across grid cells could overestimate the total number of unknown species if unknown species occur in more than one assemblage. The 220km and 440km spatial resolutions may therefore slightly underestimate percent of total discoveries, but we retained these resolutions for the visual detail they offered. All assemblage-level estimates are available through Data S3 files.

### Country-level predictions

We used the assemblage-level predictions to compute country-wide estimates of *UnknownSR* and *PropUnknown* at each spatial resolution. For each grid cell, we quantified the proportion of landcover that overlapped with countries (a same grid cell could be assigned to multiple countries, but the proportion of landcover may differ). Country selected grid cells had their values of *UnknownSR* and *PropUnknown* weighted by the respective proportion of country landcover. Given the same spatial resolution, the sum of *UnknownSR* returns the same value when computed across either countries or assemblages. We summed the *UnknownSR* across countries to get the number of unknown species per country, and averaged the *PropUnknown* to obtain the country wide proportion of unknown species. The country-level predictions were also standardized in the same way we did for taxa and assemblages. We divided the per country *UnknownSR* by the global number of *UnknownSR* to get the estimated percent of total discoveries, and standardized *PropUnknown* to vary between 0 and 1, with 1 indicating the country with the highest proportion of unknown species. Country boundaries followed the Global Administrative Units database, version 1.6 ^92^. All country-level estimates are available through Data S4 files.

## ACKNOWLEDGMENTS

We are grateful to S. Meiri, D.S. Rinnan, G. Reygondeau, N. Upham, M. Costello, D. Wake, and Joaquín Hortal for providing helpful comments on the research or manuscript drafts. We thank C. Haddad, L.C. Márquez, G. Singh, and A.F. Meyer for providing pictures of the example species in Figure 1. This work was produced, in part, with the support of the National Geographic Society. W.J. also acknowledges support from the E.O. Wilson Biodiversity Foundation and Half-Earth Project, NSF grant DEB-1441737 and NASA Grants 80NSSC17K0282 and 80NSSC18K0435.

## AUTHOR CONTRIBUTIONS

MRM and WJ conceived the study; MRM analysed the data, MRM and WJ developed the figures, MRM and WJ wrote the text.

## DATA AVAILABILITY

All datasets of species distributions used in the analyses are available at Map of Life (https://mol.org). R-scripts will be made publicly available on publication.

## SUPPLEMENTARY INFORMATION

### SUPPLEMENTARY METHODS

We performed model validations to verify if our model framework were able to satisfactorily predict species discoveries based on random species subsets left out of the model training. We also performed extensive sensitivity analysis to assess limitations of our approach to three issues detailed in the following.

#### Model validation

We used a four-fold cross-validation approach to examine the predictive accuracy of our models. For each vertebrate group, the species were randomly partitioned into four equal parts, with three of those used as the training-fold and the fourth as validation-fold. The cross-validation process was repeated four times, with each of the four-fold subsamples used once as the validation data. For each cross-validation round, we performed the model averaging approach using the 11 predictor variables and obtained for each species, the weighted average value of the (i) discovery probability in year 2014, and (ii) estimated description date.

For each training-fold, we obtained the coefficients of all possible AFT models (2,047 models covering all predictor combinations), as well as their BIC weights (wBIC). We used the model coefficients of each training-fold to predict the description dates of the 25% of species left out of the model fitting (validation-fold). These predictions were obtained for each one of the 2,047 AFT models, in each training-fold. For each species, we then calculated the weighted average of the predicted description dates using wBIC of each model as relative weights. These species-level predictions of the description date were therefore independent from the model fitting.

At this point, we proceeded with model validation at three different levels: species, taxon, assemblage.

##### 1) Species-level validation

We assessed the accuracy of the predictions of description dates from the average weighted AFT model with three different statistics. (i) The Spearman correlation measured our ability to correctly rank species according to their discovery year. (ii) The slope of the linear regression between observed and predicted description dates assessed under- or overestimation of absolute values. (iii) The normalized root mean square error (NRMSE) measured the divergence of predictions from observations ^1^. NRMSE is given in percentage, where lower values indicate less residual variance 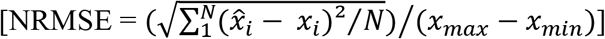. These three statistics were computed for each the four vertebrate groups. Computations were performed in R using the ‘hydroGOF’ ^2^ and ‘stats’ ^3^ packages.

##### 2) Taxon-level validation

For each cross-validation round, we calculated taxon-level estimates of *UnknownSR* as explained above. We averaged the outputs of the four cross-validation rounds to get the estimated *UnknownSR* (hereafter, estimated discoveries). Using the 25% of the species left out of the model, we obtained the observed number of unknown species (observed discoveries) per taxon, that is, the species richness per taxon based on the validation fold.

To evaluate predictive accuracy, we calculated three different statistics using the observed and estimated discoveries per taxon. (i) The Spearman correlation measured our ability to correctly rank taxonomic genera, families, and orders with respect to the number of unknown species. (ii) The slope of the linear regression between observed and estimated discoveries was used to check for under- or overestimation of the estimated discoveries. (iii) The normalized root mean square error (NRMSE). These three statistics were computed for each taxonomic rank (family, order) of each of the four vertebrate groups. Computations were performed in R using the ‘hydroGOF’ ^2^ and ‘stats’ ^3^ packages.

##### 3) Assemblage-level validation

This validation followed that conducted at the taxon level. Here, the outputs of the four cross-validation rounds were averaged to get estimated *UnknownSR* (estimated discoveries) per assemblage grid cell. We used the 25% of species in the validation fold to extract the observed number of unknown species (observed discoveries) per assemblage grid cell.

The relationship between observed and estimated discoveries per grid cell was assessed through three different statistics: (i) Spearman correlation, (ii) slope of the linear regression, and the (iii) normalized root mean square error (NRMSE). These three statistics were computed for each subsampling level (using 1, 5, 10, 50, 100, 200 and 500 grid cell occurrences per species) and the observed data (all grid cell occurrences per species), at the three spatial resolutions (grain sizes 220, 440, and 880 km), and for each vertebrate group. Computations were performed in R using the ‘hydroGOF’ ^2^ and ‘stats’ ^3^ packages.

##### 4) Country-level validation

Per country results are aggregates of the assemblage-level predictions, and are represented by the assemblage-level validation.

#### Model limitations

It is important to note that the estimated time-to-event functions derived from the AFT models may overestimate the discovery probability if the empirical time-to-event functions do not yet approach asymptote ^4^. Consequently, the estimated proportion of known species, *PropKnown*, for species in a taxon or assemblage will also be overestimated, ultimately leading to more conservative estimates of the *PropUnknown* and *UnknownSR*. We acknowledge that the discovery curve of amphibians and reptiles have not yet approach asymptote, as evidenced by the cumulative number of species description over time (Fig. 1, main text). That said, our model framework is subject to three major limitations.

First, time-to-event data are often incompletely observed, in which case the data may be considered censored and/or truncated. Broadly speaking, truncation is related to the study design and is further divided in two types. Left truncation arises from the specification of a minimum entry time for the sampling units. If the event occurs before the minimum entry time, those sampling units will never enter the study. Right truncation occurs when sampling units are only observable if they have experienced the event ^5^. Our species data shows this latter feature. The sampling condition imposed to our data may affect our estimates of discovery probability.

Second, over the temporal scale considered in this study (almost 260 years), taxonomists have dealt with different issues to describe new species. For example, earlier naturalists crossed oceans on ships to find unknown taxa in regions previously considered highly remote. In contrast, modern taxonomists have been required to use multiple tools to accumulate more evidences to provide highly detailed descriptions of new species ^6^. Although these different technological contexts are important to the discovery process, they are difficult to measure and incorporate into our models. Both predictors and model performance may vary through time. In using only species described more recently as input data, we might get different results.

And third, our model validation procedure is subject to two mathematical constraints. The number of estimated discoveries is affected by the size of the training-fold. In adding more species to the training-fold, we tend to increase the known richness per taxon or assemblage and consequently, the estimated unknown richness (*UnknownSR*). Moreover, the number of observed discoveries per taxon or assemblage is dependent on the size of the validation-fold; that is, the number of species left out of the model training. In increasing the size of validation-fold we tend to obtain higher values of observed discoveries, with the opposite occurring if we decrease the size of validation-fold. Therefore, the size of validation- and training-folds may affect how we rank taxa and regions according to their potential for future species discovery.

We investigate these three limitations futher below.

##### 1) Robustness to sampling condition

The right truncation feature creates a sampling condition, ***S***_***i***_ ≤ ***T***_***max***_, where ***S***_***i***_ is the taxonomic age of species *i* (i.e., the number of years since 1758), and ***T***_***max***_ equals the temporal range of this study (2014 – 1758 = 256 years). Only species with the taxonomic age ≤ 256 entered the study. Underlying this sampling condition is the assumption that the chance of an event at the time ***T > Tmax*** is zero [***P(T*** >***T***_***max***_) = ***0***]. This condition may be especially relevant if ***T***_***max***_ is too short, which could lead to an underrepresentation of recent-described species. Consequently, the importance of species-level attributes may depart from their true effect, leading to biased estimates of discovery probability.

Although the theoretical background to deal with incompletely observed time-to-event data has improved in the recent decades ^5^, the statistical tools available to account for right truncation are either restricted to particular time-to-event models - e.g. Cox Proportional Hazard Model ^7^ - or have somewhat limited application due to the fail in estimator convergence ^8^. Herein, we assessed the robustness of our results to the violation of this sampling condition through a sensitivity analysis. We created 10 data subsets containing increasing levels of right truncation. To do so, we successively discarded *x*% of the most recent-described species, with *x* varying from 0 to 50, at intervals of 5%. For each data subset, we repeated the model averaging framework and registered the average weighted coefficients of each predictor variable, as well as the estimated discovery probability of each species.

This sensitivity analysis is intended to answer three questions. First, how the effect size of predictors varies when the AFT models are trained with datasets holding higher levels of right-truncation? Second, how the changes in effect size (if any) affect the estimation of species’ discovery probability? To answer this latter question is important to compare the discovery probabilities for the same set of species. For this purpose, we used the oldest half of the known species, since these species were included in all data subsets. And third, what kind of the species-level attributes are expected for species yet to be described, and how those attributes would affect the estimation of discovery probability?

##### 2) Influence of the time period of species discovery

To assess the differential influence of early species discoveries in our model performance, we performed a sensitivity analysis by successively discarding previously described species. Our goal was i) to evaluate the variation of predictors across time and ii) to identify the period that offered strongest model performance for prediction. In addition to the full time period of this study (1759-2014), we defined other 22 time periods covering the interval from *d* to 2014, where *d* is a decade from 1760 to 1970 (e.g. 1760-2014, 1770-2014, …, 1970-2014). We then filtered our species dataset to include only those taxa described within each time period and repeated the model validation framework at the level of species, taxa, and assemblages. We note that with increasing *d* the sample sizes available in the datasets decreased (Table S3).

For the sensitivity analysis at the species-level, we investigated the relationship between observed and predicted description dates. At the level of taxa and assemblages, the sensitivity analyses dealt with the association between observed and estimated discoveries per taxonomic rank and assemblage grid cell, respectively. We obtained three statistics of model evaluation for the relationship of interest in each time period: (i) Spearman r, (ii) regression slope, and (iii) NRMSE. After identifying which time period returned the best model performance (see Supplementary Text section), we ran an additional analysis using the dataset for the best time period to obtain the species-level discovery metrics used to characterize discovery trends per taxon and per assemblage.

##### 3) Influence of the size of cross validation partitions

To elucidate if the mathematical constraints of our model validation affected our results, we repeated the procedure using four different sizes of validation- and training-folds. More specifically, we included 25, 50, 75, and 90% of randomly selected species in the training-fold, while keeping each respective complement (75, 50, 25, and 10% of species) in the validation-fold. We then repeated the model validation procedure at the level of taxa and assemblages, and registered three different statistics to assess the relationship between observed and estimated discoveries: (i) Spearman correlation, (ii) slope of the linear regression, and the (iii) normalized root mean square error (NRMSE). This sensitivity analysis was also repeated across the different time periods of species discovery discussed above. Computations were performed in R using the ‘hydroGOF’ ^2^ and ‘stats’ ^3^ packages.

### SUPPLEMENTARY RESULTS

#### Model Limitations and Sensitivity Analyses

##### 1) Robustness to sampling condition

We found low variation in the effect size of model coefficients if up to 30% of more recently described species were discarded before model computations (Extended Data, Fig. S1). The increasing of right-truncation decreased the effect size of model coefficients. Only a few covariates changed the effect direction with increasing levels of right-truncation. The model coefficients were nearly invariant for birds.

The variation in model coefficients is expected to influence the estimated discovery probability. In increasing the level of the right-truncation of the species dataset, we obtained higher values of discovery probability relative to those estimated from more complete datasets (Extended Data, Fig. S2). Such changes were evident mostly for amphibians and reptiles, and they were virtually nonexistent for mammals and especially birds.

Among the most consistent predictors affecting the discovery probabilities there were the (i) geographic range size, (ii) taxonomic activity per biome, and (iii) species body size. The frequency distribution of these predictor variables (Extended Data, Figs S3–S6) confirms the well-known trend of recent-described species to show narrower geographic ranges, smaller body sizes, and be described by more taxonomists relative to species described long ago ^9–11^.

It is worth noting that this sensitivity analysis indirectly considered the modelling of discovery probability for species datasets with different ending dates, in an opposite way to the analysis on the influence of the time period of species discovery (next subtopic). Here, species datasets always started in 1759 but ended at different dates, according to the percentange of recently described species discarded before computations. Thus, the ending dates were not necessarily equal for a same percentange of discarded species. For instance, the first half of the currently known amphibian species were described by 1972, whereas 50% of the bird diversity were already known by 1845 (see histograms of description year, Figure 1).

Had we been able to incorporate species-level attribute of unknown species, we would have found higher effect size for the most important model coefficients and therefore estimated lower species discovery probabilities. Given the right-truncation nature of data used in our analysis, it is likely that our discovery metrics, the *PropUnknown* and *UnknownSR* per taxon or per assemblage, are underestimated. The proportion and number of unknown species we report should be considered as conservative estimates, particularly for amphibians and reptiles.

##### 2) Influence of the time period of species discovery

We evaluated the sensitivity of our ‘discovery metrics’ to the time period of species discovery (e.g. 1760-2014, 1770-2014, …, 1970-2014). In discarding species described long ago, most of the model coefficients decreased their effect size (Fig. 2, main text). The ability of the AFT models to correctly predict the species description dates did not improve after excluding earlier-described species, except in mammals (but at the cost of discarding more than 70% of all known mammal species; Table S3, Extended Data, Fig. S7).

At the taxon-level, the Spearman correlation between observed and estimated discoveries was roughly constant after discarding species described during the first century of the modern taxonomy (Extended Data, Fig. S8). At the assemblage level, the model performance also decreased as old described species were discarded before computations. The decline in model performance was evident across all subsampling levels used to control the overrepresentation of wide-ranging species (Extended Data, Figs S9–S12). The subsampling level of 5 occurrences per species showed the best performance in explaining the observed discoveries per grid cell, a result consistent across the all spatial resolutions and vertebrate classes.

Overall, we did not obtain better measures of estimated discoveries by removing species described long ago. We therefore used the complete dataset (species described from 1759 to 2014) to estimate the discovery probability at the species-level and to estimate the number of unknown species (*UnknownSR*) at the taxon- and assemblage-level. Overall, we found a strong association with description year in extracting the predicted year of discovery at the threshold of 0.5 of discovery probability (Spearman *r* = 0.65–0.81 for all groups, see Extended Data, Fig. S13).

##### 3) Influence of the size of cross-validation partitions

Across taxonomic ranks, we found similar values of Spearman correlation between observed and estimated discoveries, regardless of the size of the species dataset used in the model training and mapping procedure (Extended Data, Fig. S8). The size of cross-validation partitions did not influence model performance when using different time periods of species discovery either (Extended Data, Fig. S8). The extraction of *PropUnknown* based on small sample sizes (i.e., using 25% of species in the model training) were less able to properly characterize discovery patterns at the taxon-level relative to large sample sizes (including 50, 75, 90% of species in the model training).

The regression slope between observed and estimated discoveries per taxon tended to decrease when assessed at higher taxonomic ranks and for training-folds containing more species (Extended Data, Fig. S8). The estimated discoveries underestimated the observed discoveries across all taxonomic ranks, although such underestimation were less pronounced when using 90% of species in the model training (Table S4). After standardizing *UnknownSR* to percent of total discoveries, we observed similar model performance for all sizes of the cross-validation partitions (Extended Data, Fig. S14).

At the assemblage-level, the relationship between observed and estimated discoveries showed similar values of Spearman correlation, regardless of the size of cross-validation partitions (Extended Data, Figs S9–S12). Once again, subsampling 5 grid cell occurrences per species resulted in estimated discoveries that better predicted the observed discoveries per grid cell, regardless of the size of cross validation partitions. The discrepancy between the absolute number of observed and estimated discoveries per grid cell decreased when species assemblages were defined at either coarser spatial resolutions or based in training-folds including higher proportion of randomly selected species (Table S4). After standardizing *UnknownSR* to percent of total discoveries, the relationship between estimated and observed discoveries per grid cell was also constant for all sizes of the cross-validation partitions (Extended Data, Fig. S15).

Overall, the estimated discoveries we obtained, either at the taxon- or assemblage-level analyses, underestimated the observed discoveries. The underestimation was higher when the discovery metrics were computed for models including fewer species (Table S4). We therefore recommend caution in interpreting the absolute values of the estimated number of unknown species (*UnknownSR*) per taxon or per grid cell. We reinforce the strong monotonicity between the observed and estimated discoveries, which ultimately support our ability to rank taxa and regions according to their potential for the discovery of new species. We do not advise using this approach to update global numbers of unknown species, unless extensive cross validation reveal absence of under- or overestimation of *UnknownSR*.

##### 4) Species authority name assumptions

Our time-to-description analysis is based on original species authority names and uses the year in which a given binomial was originally proposed. Authority name description years reveal patterns of species discovery over time ^4,9–15^, but do not account for the complicated taxonomic history of synonymizations and revalidations associated with a binomial name. A species that was recently revalidated still holds the same authority name – and therefore description year – as when it was first recognized as unique taxon. Therefore, our findings concern factors affecting the year when a species was first discovered, and not if and when a revalidation occurred.

It is worth noting that a species description is a scientific hypothesis ^16^, which can be revisited if more data become available, as often illustrated in integrative taxonomy studies ^17^. It is not uncommon for broad studies such as taxonomic reviews to split a previously described species into multiple species, and in some cases, to resurrect synonyms or elevate subspecies to species rank. Although such ‘splitting’ might sometimes be viewed as undesirable ^18,19^, when driven by scientific insights is a vital part of the taxonomic knowledge evolution, as taxonomies are not static over time ^16,20^. Among 149 integrative taxonomy studies recently published in vertebrates (including fish), 40% consider all species as valid without changes, 31% pointed out at least one undescribed species but did not formally describe it, and 30% described at least one new species ^6^. Thus, at least among vertebrates, the identification of new lineages (if any) is not necessarily followed by the proposition of new names.

Many firsthand species discoveries, i.e. species descriptions that propose new authority binomials, are reported for vertebrates every year. For example, more than 85% of amphibian species described between 1992-2003 resulted from newly proposed names, with less than 15% of descriptions concerning elevation of subspecies to species or revalidation of synonymies ^21^. In reptiles, 79% of species descriptions between 1992-2017 were published outside revisionary taxonomic studies ^22^. Newly proposed binomial names are also a large part of recently described mammals. Since 2005, 1,251 new mammal species have been recognized as valid, with 42% of them comprising firsthand discoveries and 58% consisting of resurrection of synomymies or elevation of subspecies to species rank ^23^. Altogether, these estimates illustrate the size of the taxonomic enterprise ahead. Concerns might go beyond differentiating firsthand discoveries from species revalidations, questioning the validity of species descriptions under the application of different species concepts ^18^. To date, reflecting the ongoing debate about species concepts in biology ^24^, comprehensive taxonomic databases that standardize global species lists according to a single species concept remain out of reach.

## SUPPLEMENTARY TABLES

**Table S1.** Taxon-specific allometric equations used to convert straight carapace length (SCL) of chelonians into body mass. Sample size used to create the equations, number of species covered by the samples, and summary statistics of each allometric equation are shown. Allometric equation: Log(BodyMass) = Intercept + Coef × Log(SCL).

| Taxa | Sample Size | Number of species | Intercept | Coefficient | R<sup>2</sup> |
| --- | --- | --- | --- | --- | --- |
| Chelidae | 52 | 42 | -3.915 | 2.991 | 0.947 |
| Emydidae | 82 | 57 | -3.420 | 2.814 | 0.950 |
| Kinosternidae | 28 | 27 | -3.235 | 2.715 | 0.913 |
| Podocnemididae | 14 | 8 | -3.757 | 2.930 | 0.952 |
| Testudinidae | 96 | 52 | -3.473 | 2.885 | 0.956 |
| Cryptodira<sup>*</sup> | 326 | 218 | -3.389 | 2.832 | 0.928 |
| Pleurodira<sup>*</sup> | 68 | 49 | -3.922 | 2.995 | 0.967 |

<sup>\*</sup> Only used when a family-level equation was not available.

**Table S2.** Variance Inflation Factor (VIF) for species-level attributes of terrestrial vertebrates using the full dataset (species described from 1759 to 2014). VIF measures the multicollinearity of variables included in a model, and it varies from 1 (no multicollinearity) to +Inf. VIF values > 10 reflect high multicollinearity ^25^.

| **Variable** | **Amphibians** | **Reptiles** | **Mammals** | **Birds** |
| --- | --- | --- | --- | --- |
| Annual mean temperature | 2.695 | 2.364 | 2.590 | 2.615 |
| Annual precipitation | 2.661 | 2.251 | 2.424 | 2.817 |
| Body size | 1.125 | 1.192 | 1.074 | 1.072 |
| Elevation | 2.048 | 1.872 | 1.578 | 1.858 |
| Human density | 1.368 | 1.442 | 1.423 | 1.410 |
| Precipitation seasonality | 1.631 | 1.663 | 1.702 | 2.116 |
| Range size | 1.516 | 2.158 | 1.913 | 2.390 |
| Range rarity | 1.752 | 1.606 | 1.674 | 1.932 |
| Activity per bioregion | 2.155 | 2.184 | 2.454 | 1.619 |
| Activity per family | 1.922 | 1.953 | 2.311 | 1.322 |
| Temperature seasonality | 1.908 | 2.620 | 3.646 | 4.098 |

All variables were log10 transformed before computation of the Variance Inflation Factor.

**Table S3.**
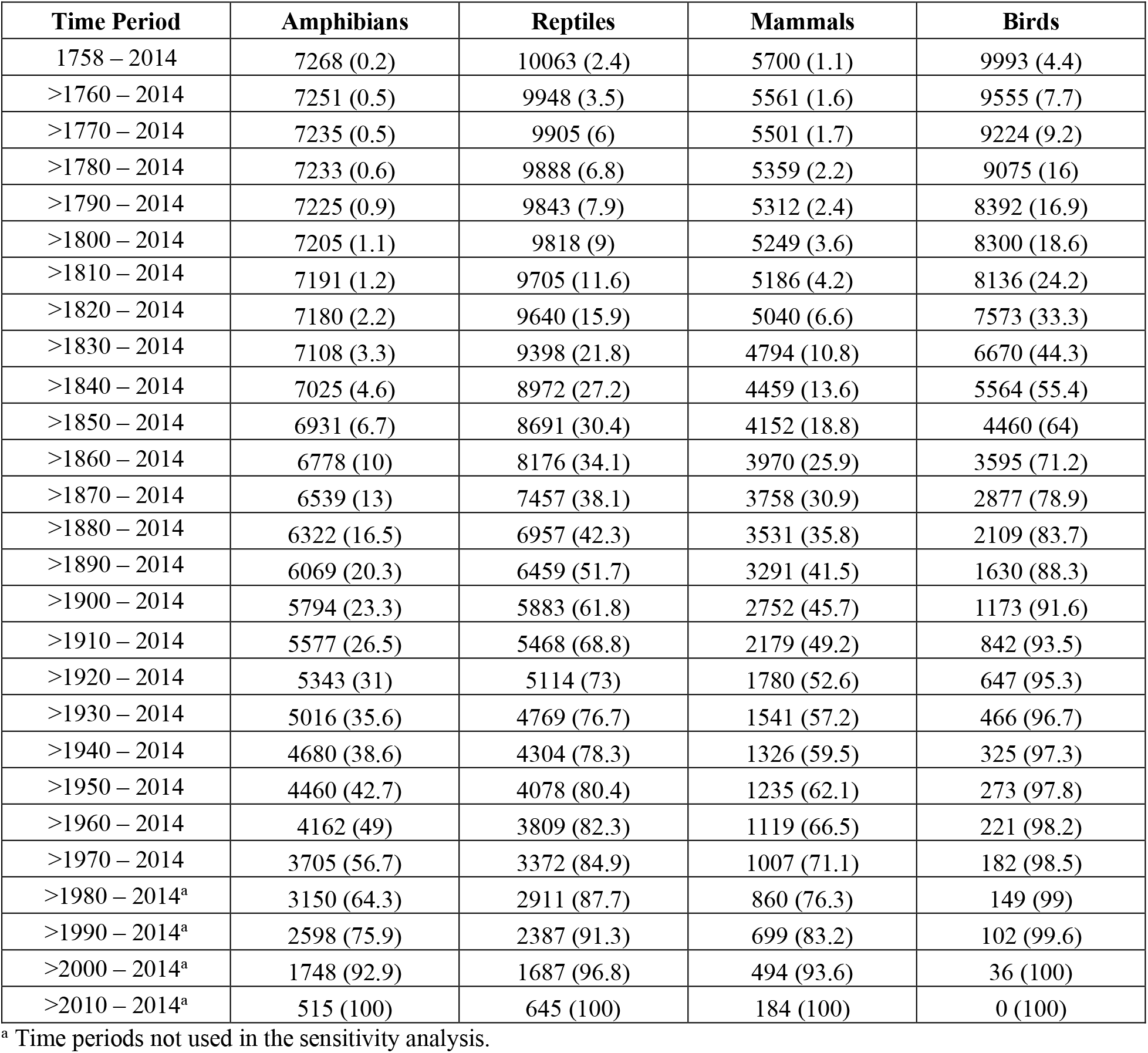
Total number of species described within the different time periods. The number between parentheses indicates the percentage of species discarded if such decades are left out of the species dataset. Only species included in our dataset were counted.

| **Time Period** | **Amphibians** | **Reptiles** | **Mammals** | **Birds** |
| --- | --- | --- | --- | --- |
| 1758 – 2014 | 7268 (0.2) | 10063 (2.4) | 5700 (1.1) | 9993 (4.4) |
| >1760 – 2014 | 7251 (0.5) | 9948 (3.5) | 5561 (1.6) | 9555 (7.7) |
| >1770 – 2014 | 7235 (0.5) | 9905 (6) | 5501 (1.7) | 9224 (9.2) |
| >1780 – 2014 | 7233 (0.6) | 9888 (6.8) | 5359 (2.2) | 9075 (16) |
| >1790 – 2014 | 7225 (0.9) | 9843 (7.9) | 5312 (2.4) | 8392 (16.9) |
| >1800 – 2014 | 7205 (1.1) | 9818 (9) | 5249 (3.6) | 8300 (18.6) |
| >1810 – 2014 | 7191 (1.2) | 9705 (11.6) | 5186 (4.2) | 8136 (24.2) |
| >1820 – 2014 | 7180 (2.2) | 9640 (15.9) | 5040 (6.6) | 7573 (33.3) |
| >1830 – 2014 | 7108 (3.3) | 9398 (21.8) | 4794 (10.8) | 6670 (44.3) |
| >1840 – 2014 | 7025 (4.6) | 8972 (27.2) | 4459 (13.6) | 5564 (55.4) |
| >1850 – 2014 | 6931 (6.7) | 8691 (30.4) | 4152 (18.8) | 4460 (64) |
| >1860 – 2014 | 6778 (10) | 8176 (34.1) | 3970 (25.9) | 3595 (71.2) |
| >1870 – 2014 | 6539 (13) | 7457 (38.1) | 3758 (30.9) | 2877 (78.9) |
| >1880 – 2014 | 6322 (16.5) | 6957 (42.3) | 3531 (35.8) | 2109 (83.7) |
| >1890 – 2014 | 6069 (20.3) | 6459 (51.7) | 3291 (41.5) | 1630 (88.3) |
| >1900 – 2014 | 5794 (23.3) | 5883 (61.8) | 2752 (45.7) | 1173 (91.6) |
| >1910 – 2014 | 5577 (26.5) | 5468 (68.8) | 2179 (49.2) | 842 (93.5) |
| >1920 – 2014 | 5343 (31) | 5114 (73) | 1780 (52.6) | 647 (95.3) |
| >1930 – 2014 | 5016 (35.6) | 4769 (76.7) | 1541 (57.2) | 466 (96.7) |
| >1940 – 2014 | 4680 (38.6) | 4304 (78.3) | 1326 (59.5) | 325 (97.3) |
| >1950 – 2014 | 4460 (42.7) | 4078 (80.4) | 1235 (62.1) | 273 (97.8) |
| >1960 – 2014 | 4162 (49) | 3809 (82.3) | 1119 (66.5) | 221 (98.2) |
| >1970 – 2014 | 3705 (56.7) | 3372 (84.9) | 1007 (71.1) | 182 (98.5) |
| >1980 – 2014<sup>a</sup> | 3150 (64.3) | 2911 (87.7) | 860 (76.3) | 149 (99) |
| >1990 – 2014<sup>a</sup> | 2598 (75.9) | 2387 (91.3) | 699 (83.2) | 102 (99.6) |
| >2000 – 2014<sup>a</sup> | 1748 (92.9) | 1687 (96.8) | 494 (93.6) | 36 (100) |
| >2010 – 2014<sup>a</sup> | 515 (100) | 645 (100) | 184 (100) | 0 (100) |

<sup>a</sup> Time periods not used in the sensitivity analysis.

**Table S4.** Validation-fold and grouping dependence of deviation from a 1:1 relationship. Values shown how many times, on median, the observed discoveries were higher than the estimated discoveries. Results shown for the complete time period of species discoveries (1759-2014). At the assemblage level, only the subsampling level of 5 occurrences per species is shown.

| Grouping level | Validation-fold size | Amphibians | Reptiles | Mammals | Birds |
| --- | --- | --- | --- | --- | --- |
| **Taxon-level** |  |  |  |  |  |
| Family | 75% | 15.922 | 17.560 | 14.072 | 27.250 |
| Family | 50% | 7.312 | 8.270 | 8.329 | 18.500 |
| Family | 25% | 3.025 | 3.275 | 3.996 | 9.000 |
| Family | 10% | 1.338 | 1.349 | 1.791 | 4.200 |
| **Family** | **Average** | **6.899** | **7.613** | **7.047** | **14.737** |
| Higher-level grouping | 75% | 28.986 | 27.605 | 30.016 | 68.500 |
| Higher-level grouping | 50% | 9.887 | 11.843 | 11.402 | 46.000 |
| Higher-level grouping | 25% | 3.423 | 4.312 | 5.172 | 20.361 |
| Higher-level grouping | 10% | 1.152 | 1.469 | 1.888 | 7.392 |
| **Higher-level grouping** | **Average** | **10.862** | **11.307** | **12.120** | **35.563** |
| **Assemblage-level** |  |  |  |  |  |
| 220 km | 75% | 3.894 | 6.396 | 3.695 | 4.736 |
| 220 km | 50% | 3.068 | 4.147 | 2.957 | 3.662 |
| 220 km | 25% | 2.271 | 2.458 | 2.313 | 2.683 |
| 220 km | 10% | 2.022 | 2.017 | 2.051 | 2.211 |
| **220 km** | **Average** | **2.814** | **3.754** | **2.754** | **3.323** |
| 440 km | 75% | 9.430 | 14.658 | 9.687 | 11.627 |
| 440 km | 50% | 5.882 | 7.795 | 6.412 | 8.120 |
| 440 km | 25% | 3.097 | 3.524 | 3.487 | 4.577 |
| 440 km | 10% | 2.079 | 2.069 | 2.241 | 2.788 |
| **440 km** | **Average** | **5.112** | **7.012** | **5.457** | **6.778** |
| 880 km | 75% | 18.794 | 25.063 | 24.776 | 29.200 |
| 880 km | 50% | 9.964 | 11.985 | 14.881 | 19.592 |
| 880 km | 25% | 4.269 | 4.526 | 6.744 | 10.296 |
| 880 km | 10% | 2.211 | 2.229 | 2.981 | 4.751 |
| **880 km** | **Average** | **8.810** | **10.951** | **12.345** | **15.960** |

## SUPPLEMENTARY FIGURES

**Fig. S1.**
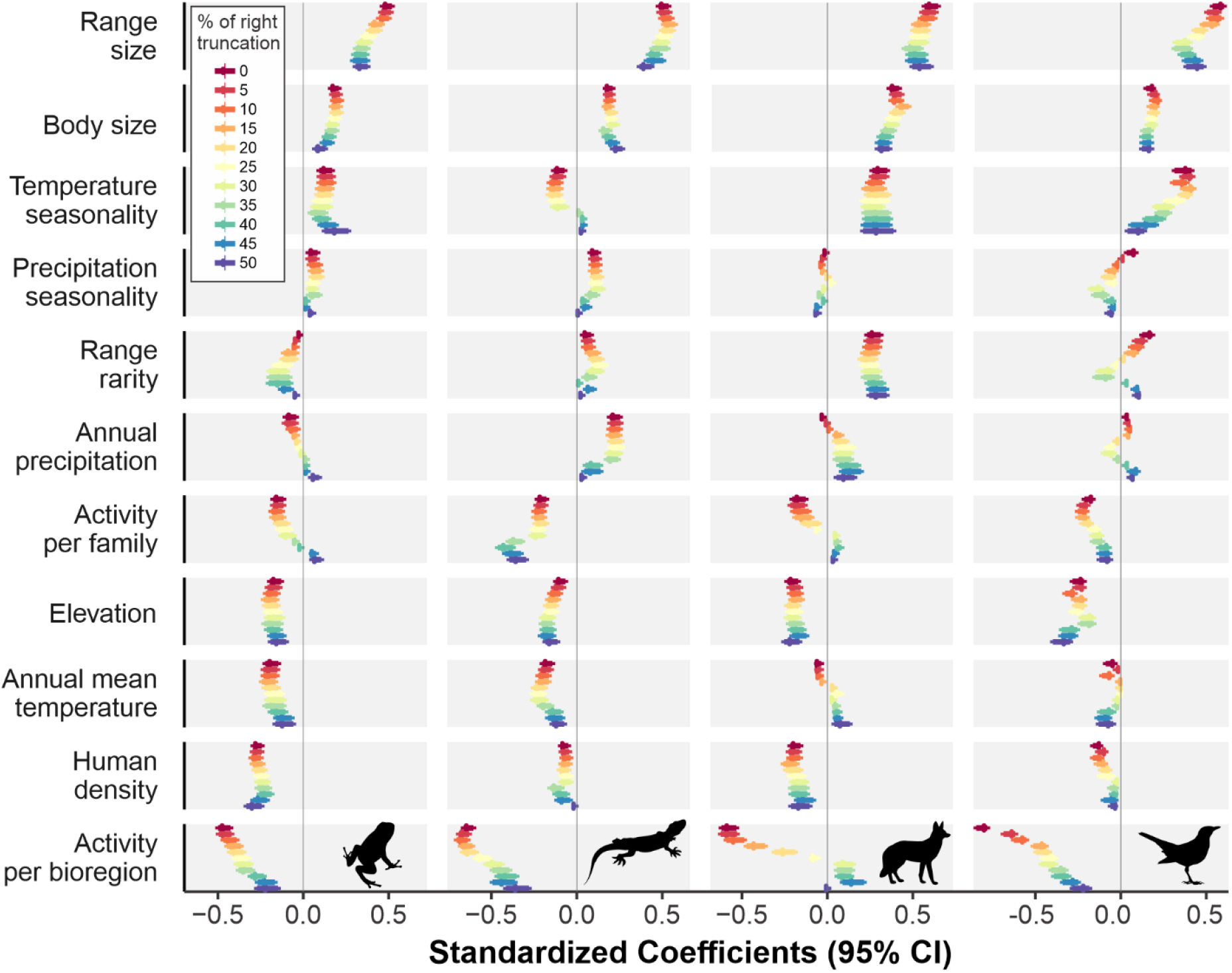
Standardized coefficients of the average weighted accelerated failure time (AFT) model computed for species datasets with increasing levels of right-truncation. Line colours indicate the percentage of recent-described species left out of the model (recent-described species were successively discarded). The horizontal bars denote the 95% confidence intervals around each coefficient. Standardized coefficients above 0 indicate that species with high values for a given attribute had higher discovery probability (prob.) and thus were likely discovered early on. Negative standardized coefficients mean high attribute values depressed discovery probability and delayed discovery.

**Fig. S2.**
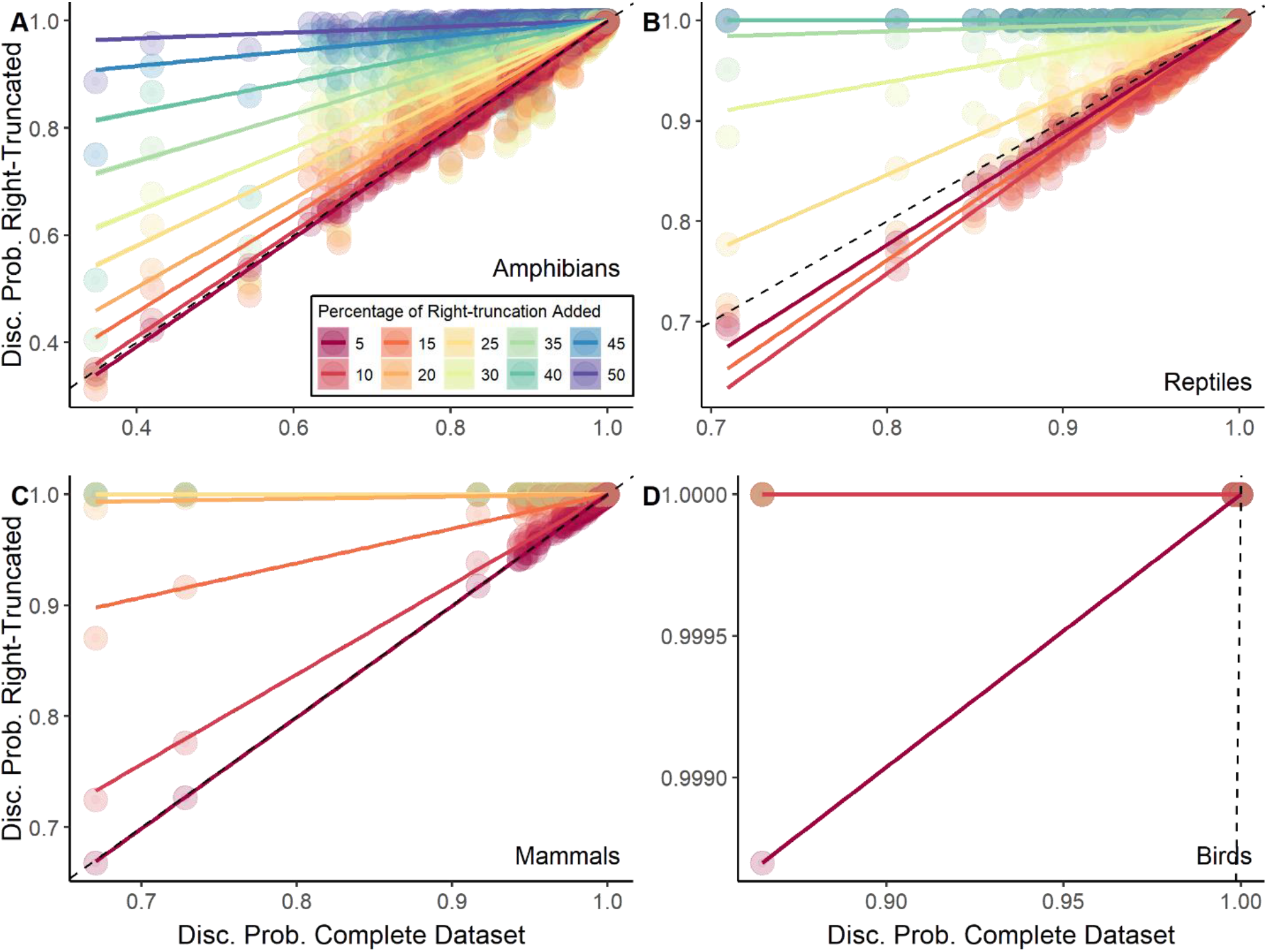
Relationship between values of discovery probability computed using the complete (x-axis) and right-truncated (y-axis) datasets. For each vertebrate group, only the oldest half of the known species is represented due to their presence in all data subsets with different levels of right-truncation. The dashed line indicates the line of equality. In decreasing the level of right-truncation (increasing completeness) of species dataset, the discovery probabilities tend to be lower.

**Fig. S3.**
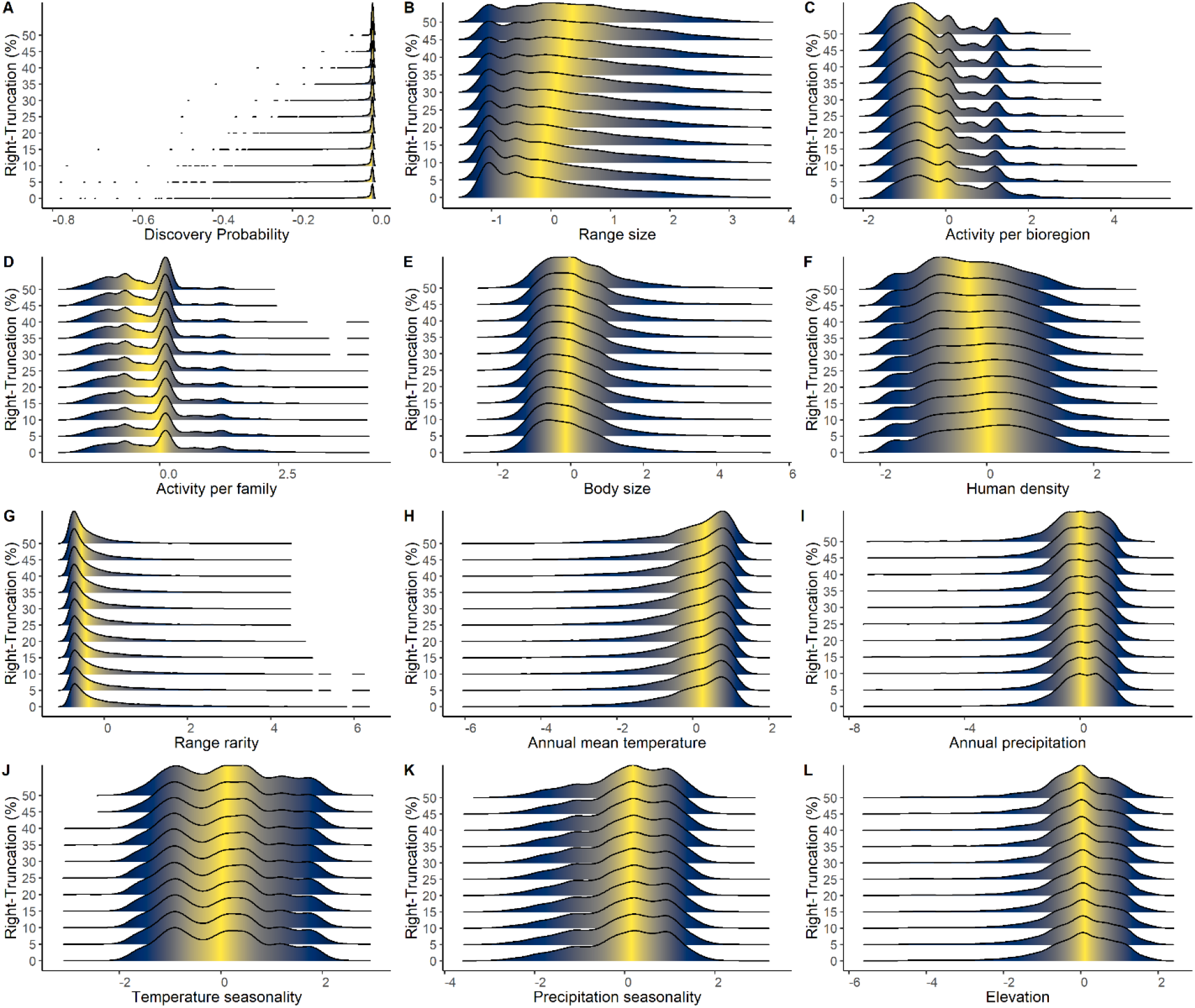
Frequency distribution of species-level attributes for data subsets with different levels of right-truncation for amphibians. (A) Discovery probability. (B) Range size = number of 110 × 110 km grid cells occupied by the species’ range. (C) Activity/bioregion = number of taxonomists per species in the bioregions in which it typically occurs at the year of the species’ description. (D) Activity/family = number of taxonomists per species in a family at the year of species’ description. (E) Body size = maximum body size. (F) Human density = Within-range human population density at the year of species’ description. (G) Range rarity = within-range endemism richness at the year of the species’ description. (H) Annual mean temperature = within-range annual mean temperature. (I) Annual precipitation = within-range annual precipitation. (J) Temperature seasonality = within-range temperature seasonality. (K) Precipitation seasonality = within-range precipitation seasonality. (L) Mean elevation = within-range mean elevation. In all plots, the variable was log_10_ transformed to increase readability. The colour gradient is centred in the median value.

**Fig. S4.**
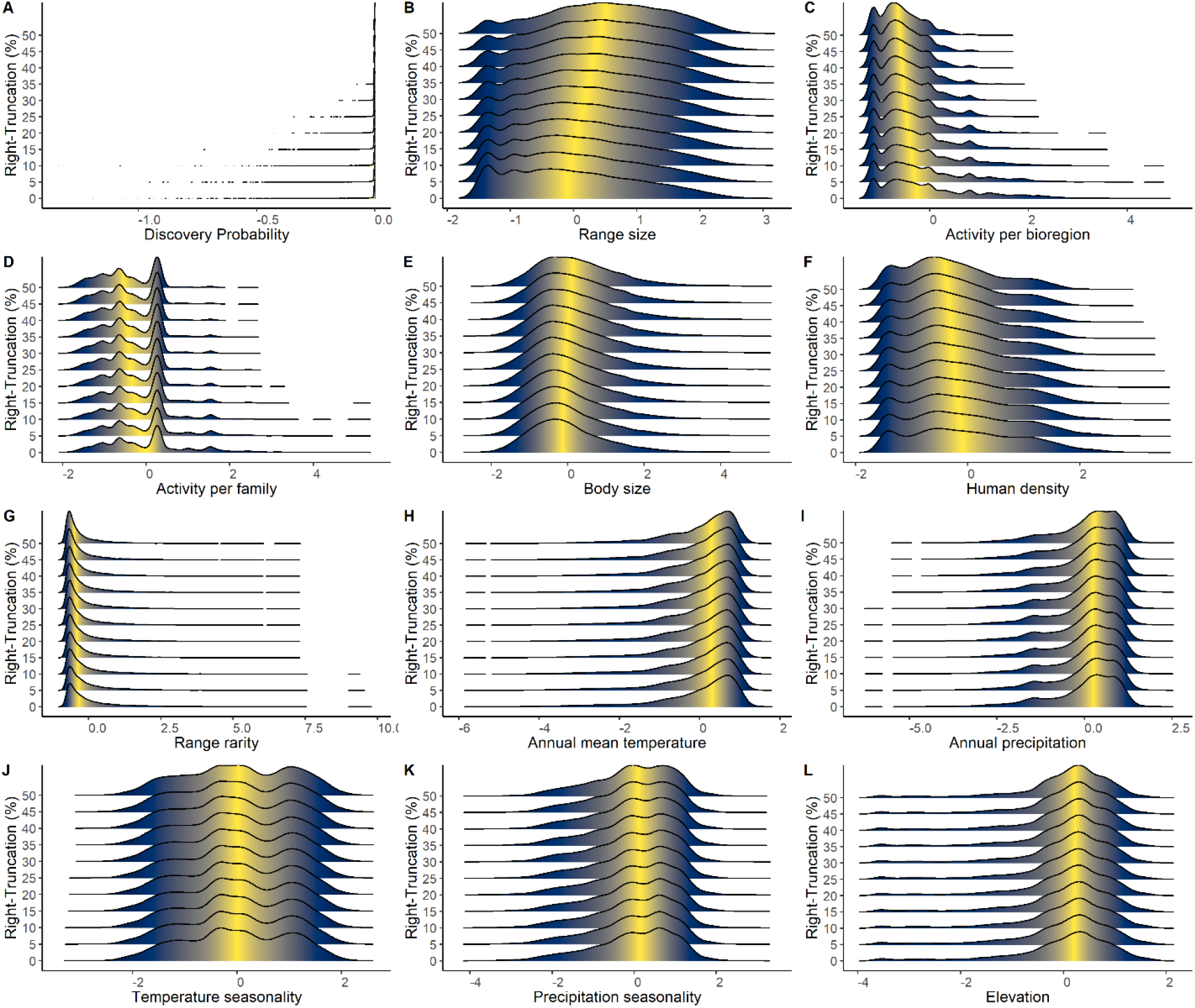
Frequency distribution of species-level attributes for data subsets with different levels of right-truncation for reptiles. (A) Discovery probability. (B) Range size = number of 110 × 110 km grid cells occupied by the species’ range. (C) Activity/bioregion = number of taxonomists per species in the bioregions in which it typically occurs at the year of the species’ description. (D) Activity/family = number of taxonomists per species in a family at the year of species’ description. (E) Body size = maximum body size. (F) Human density = Within-range human population density at the year of species’ description. (G) Range rarity = within-range endemism richness at the year of the species’ description. (H) Annual mean temperature = within-range annual mean temperature. (I) Annual precipitation = within-range annual precipitation. (J) Temperature seasonality = within-range temperature seasonality. (K) Precipitation seasonality = within-range precipitation seasonality. (L) Mean elevation = within-range mean elevation. In all plots, the variable was log_10_ transformed to increase readability. The colour gradient is centred in the median value.

**Fig. S5.**
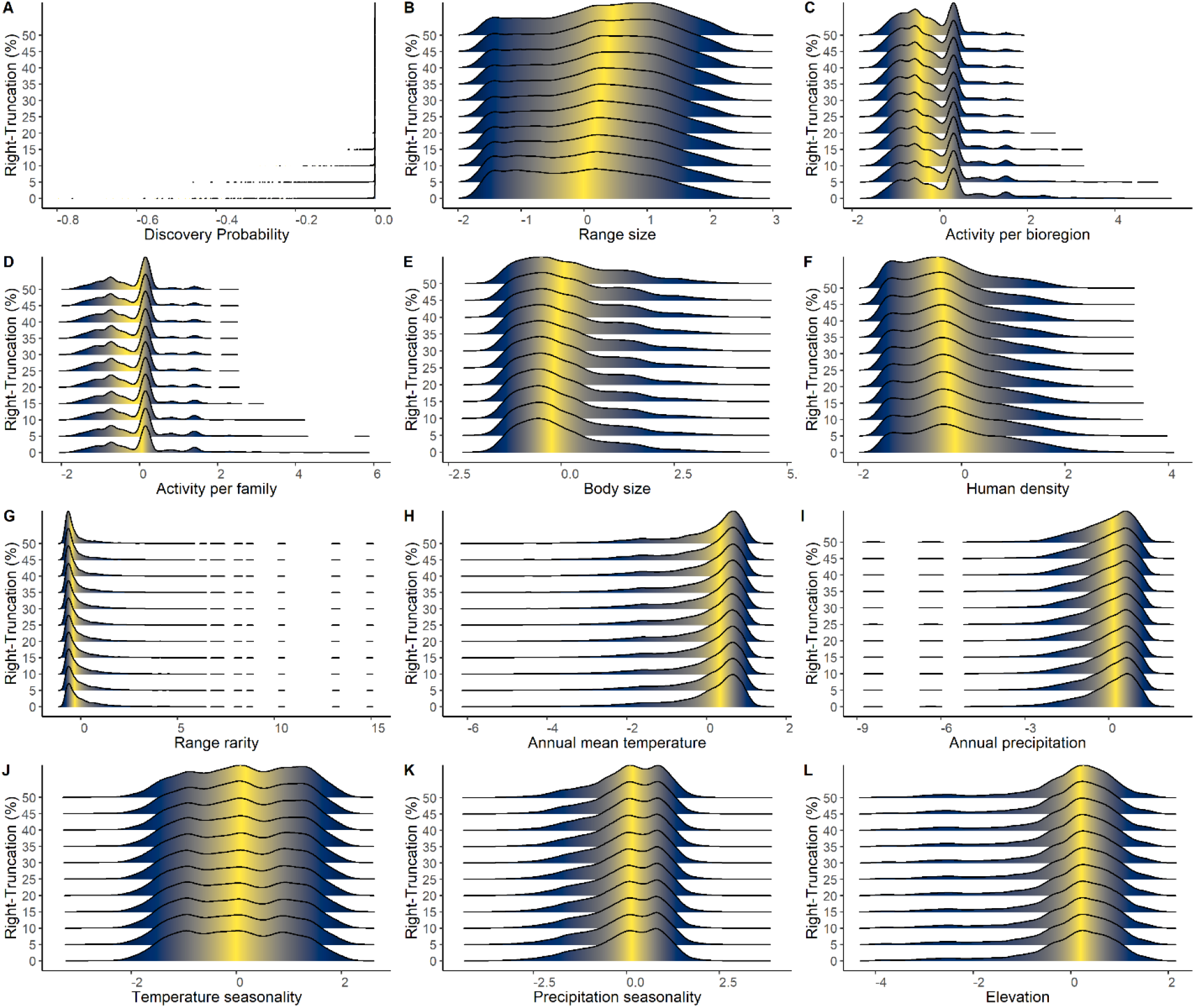
Frequency distribution of species-level attributes for data subsets with different levels of right-truncation for mammals. (A) Discovery probability. (B) Range size = number of 110 × 110 km grid cells occupied by the species’ range. (C) Activity/bioregion = number of taxonomists per species in the bioregions in which it typically occurs at the year of the species’ description. (D) Activity/family = number of taxonomists per species in a family at the year of species’ description. (E) Body size = maximum body size. (F) Human density = Within-range human population density at the year of species’ description. (G) Range rarity = within-range endemism richness at the year of the species’ description. (H) Annual mean temperature = within-range annual mean temperature. (I) Annual precipitation = within-range annual precipitation. (J) Temperature seasonality = within-range temperature seasonality. (K) Precipitation seasonality = within-range precipitation seasonality. (L) Mean elevation = within-range mean elevation. In all plots, the variable was log_10_ transformed to increase readability. The colour gradient is centred in the median value.

**Fig. S6.**
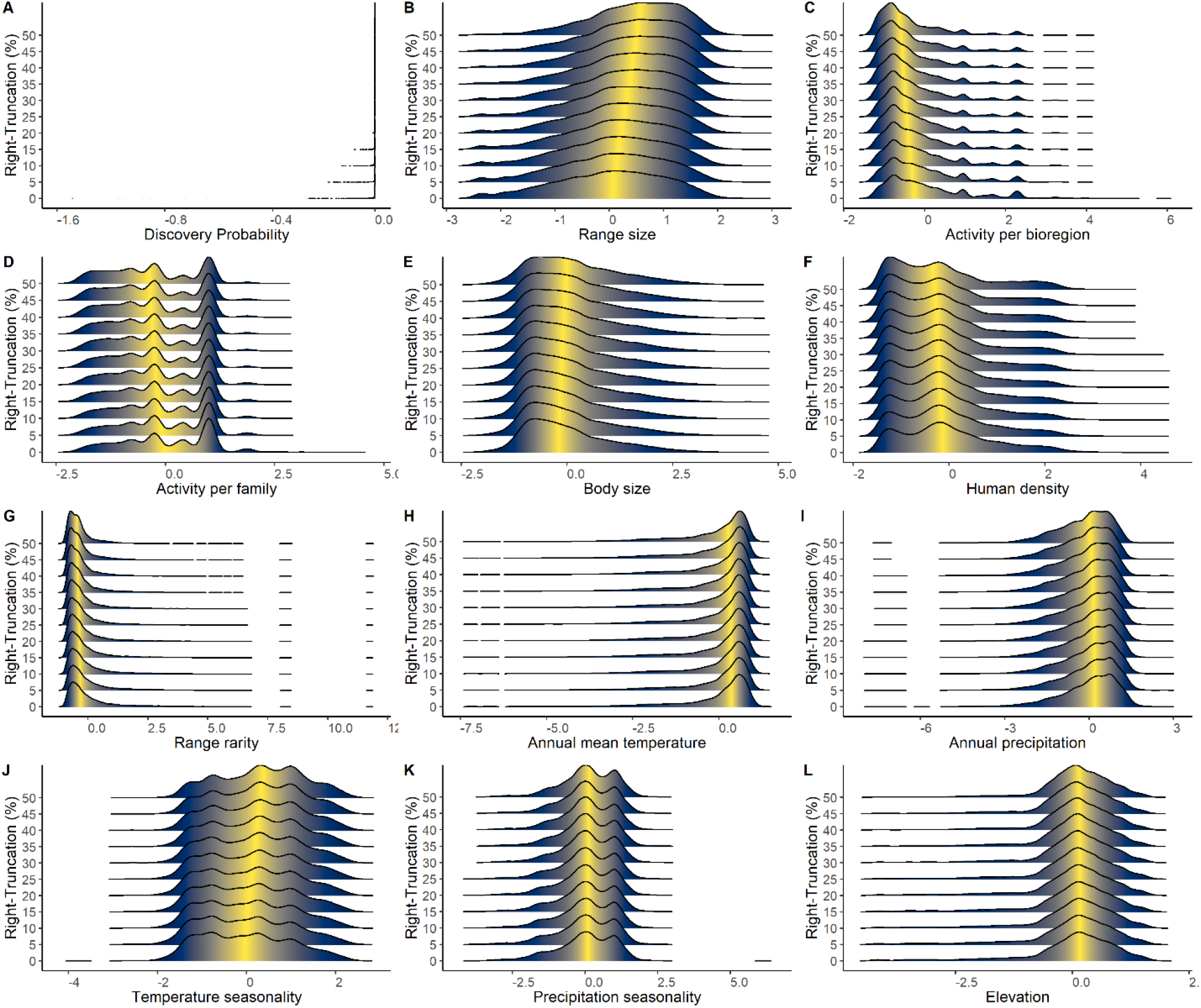
Frequency distribution of species-level attributes for data subsets with different levels of right-truncation for birds. (A) Discovery probability. (B) Range size = number of 110 × 110 km grid cells occupied by the species’ range. (C) Activity/bioregion = number of taxonomists per species in the bioregions in which it typically occurs at the year of the species’ description. (D) Activity/family = number of taxonomists per species in a family at the year of species’ description. (E) Body size = maximum body size. (F) Human density = Within-range human population density at the year of species’ description. (G) Range rarity = within-range endemism richness at the year of the species’ description. (H) Annual mean temperature = within-range annual mean temperature. (I) Annual precipitation = within-range annual precipitation. (J) Temperature seasonality = within-range temperature seasonality. (K) Precipitation seasonality = within-range precipitation seasonality. (L) Mean elevation = within-range mean elevation. In all plots, the variable was log_10_ transformed to increase readability. The colour gradient is centred in the median value.

**Fig. S7.**
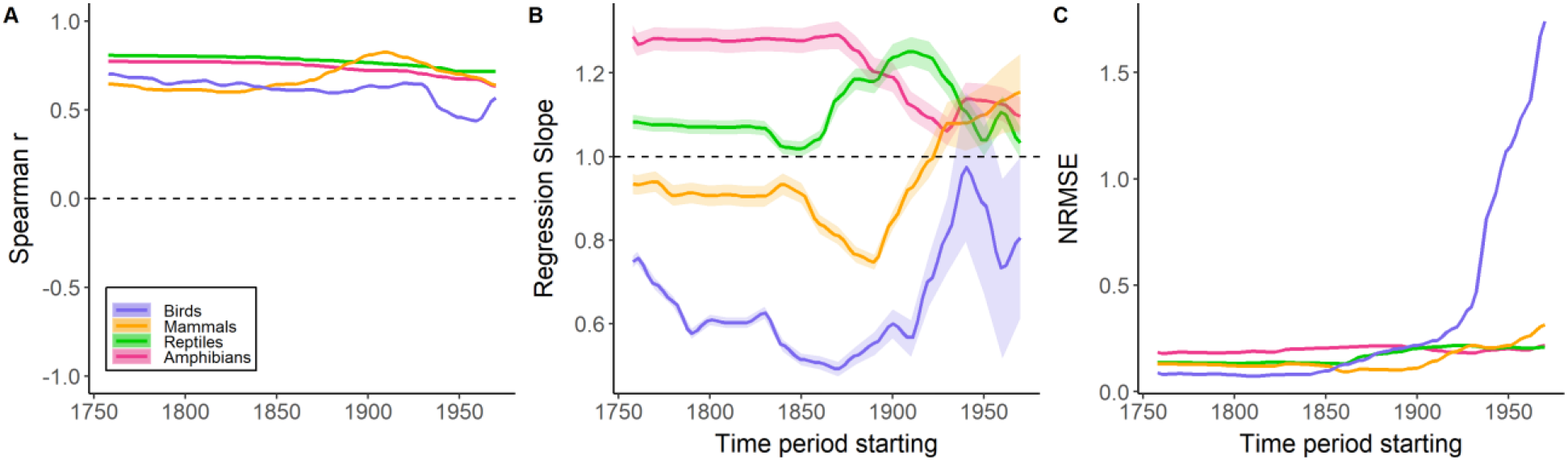
Statistic metrics derived from the sensitivity analysis at the species level. All statistics were computed between predicted and observed year of discovery across species. Results based on model trained with 75% of species and 25% of species used as holdout data. Line colours denote different vertebrate groups. (A) Spearman correlation, (B) Regression slope, (C) Normalized Root Mean Square Error (NRMSE).

**Fig. S8.**
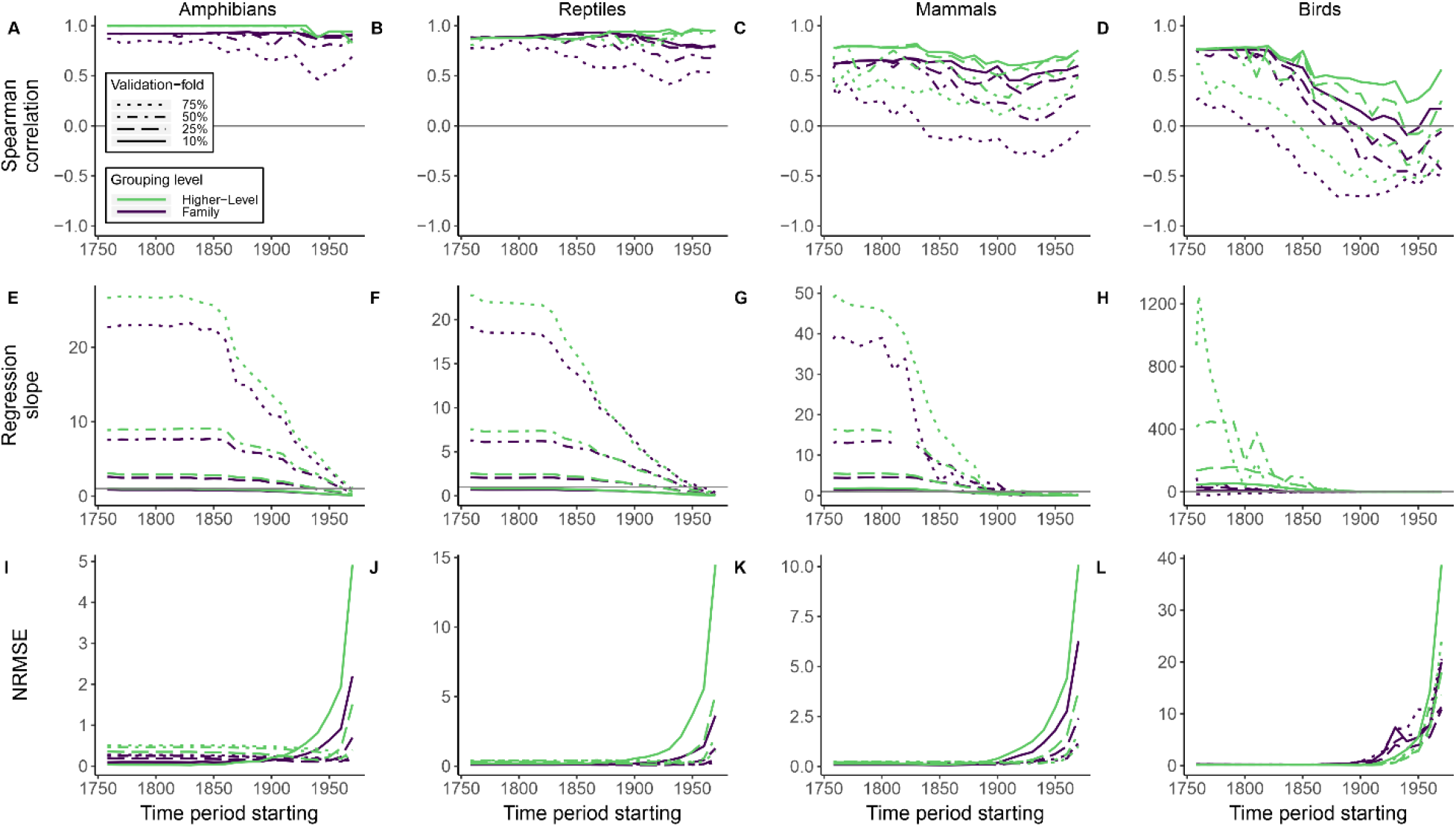
Statistic metrics derived from the sensitivity analysis at the taxon level using different sizes of validation-fold. All statistics were computed between estimated and observed discoveries across taxa. Line types denote different sizes of the validation-fold. Line colours indicate different taxonomic ranks. (A-D) Spearman correlation, (E-H) Regression slope, (I-L) Normalized Root Mean Square Error (NRMSE). The size of training-fold (25, 50, 75, 90% of species) is the complement of the respective validation-fold size (75, 50, 25, 10% of species). Confidence intervals were omitted to increase readability.

**Fig. S9.**
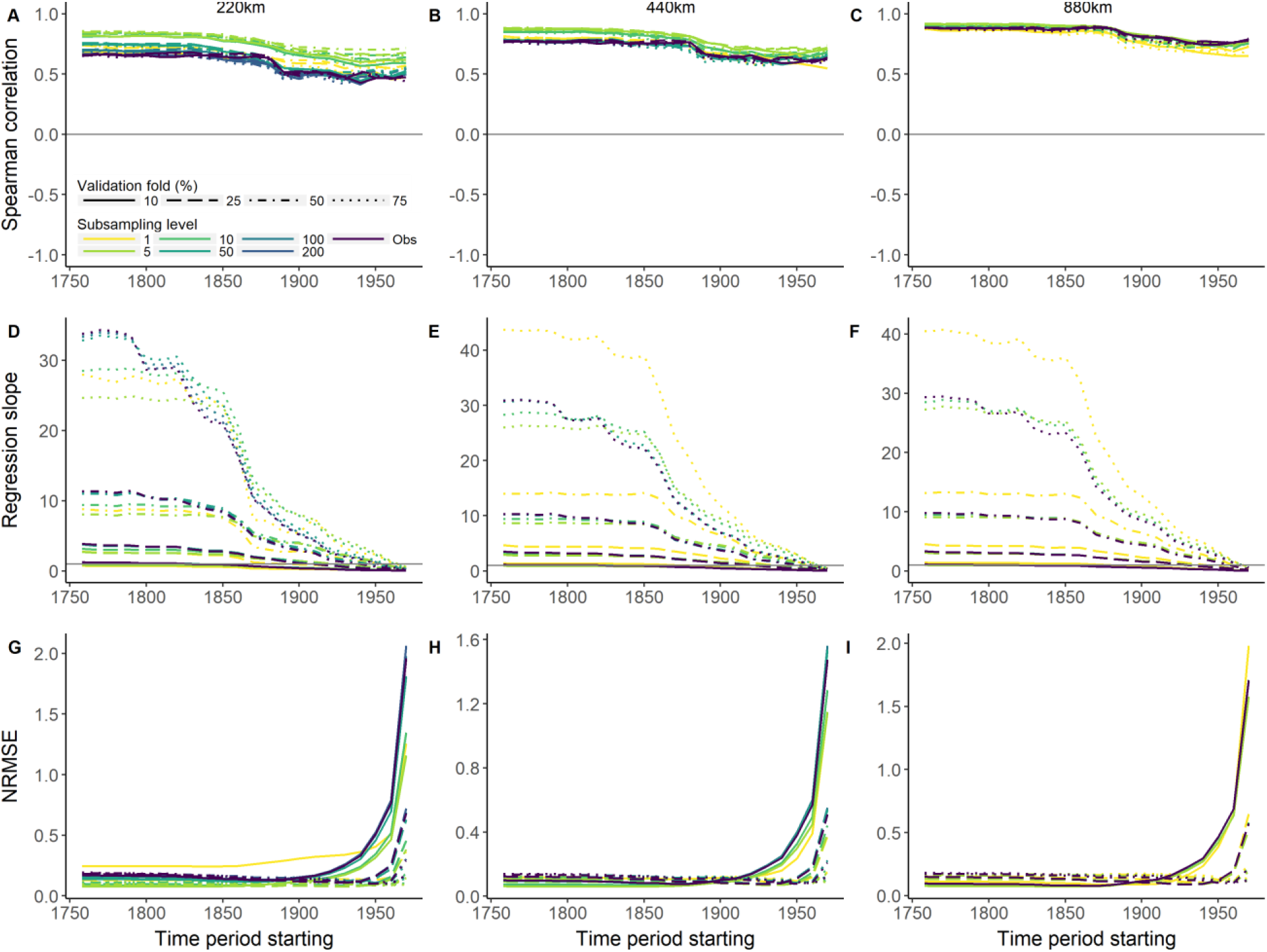
Statistic metrics derived from the sensitivity analysis using amphibian species. All statistics were computed between estimated and observed discoveries across grid cells. Line colours denote the subsampling level adopted to control overrepresentation of wide-ranging species. Panel columns refer to statistics calculated at different spatial resolutions (220, 440, 880 km). (A-C) Spearman correlation, (D-F) Regression slope, (G-I) Normalized Root Mean Square Error (NRMSE). Confidence intervals were omitted to increase readability.

**Fig. S10.**
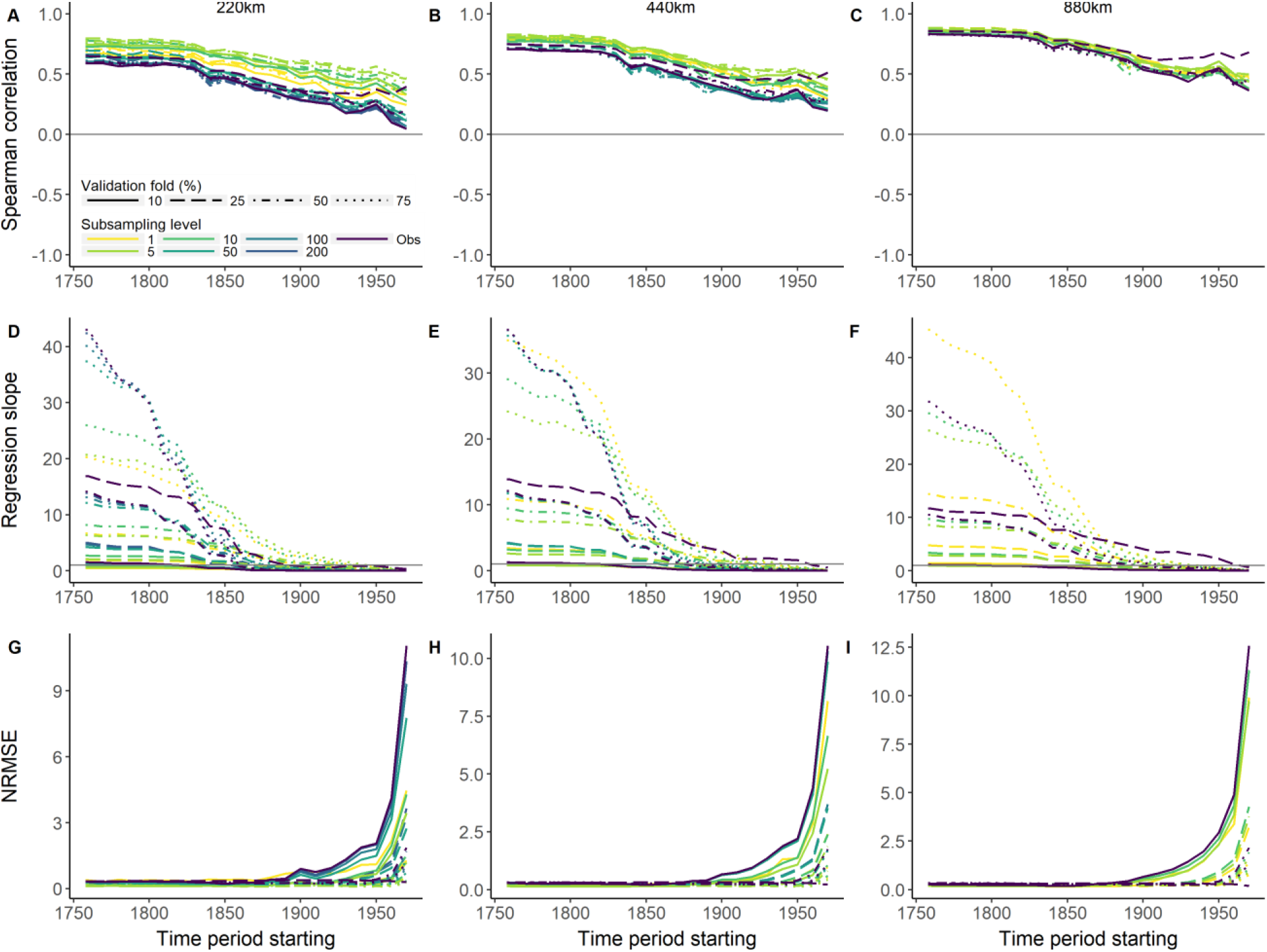
Statistic metrics derived from the sensitivity analysis using reptile species. All statistics were computed between estimated and observed species across grid cells. Line colours denote the subsampling level adopted to control overrepresentation of wide-ranging species. Panel columns refer to statistics calculated at different spatial resolutions (220, 440, 880 km). (A-C) Spearman correlation, (D-F) Regression slope, (G-I) Normalized Root Mean Square Error (NRMSE). Confidence intervals were omitted to increase readability.

**Fig. S11.**
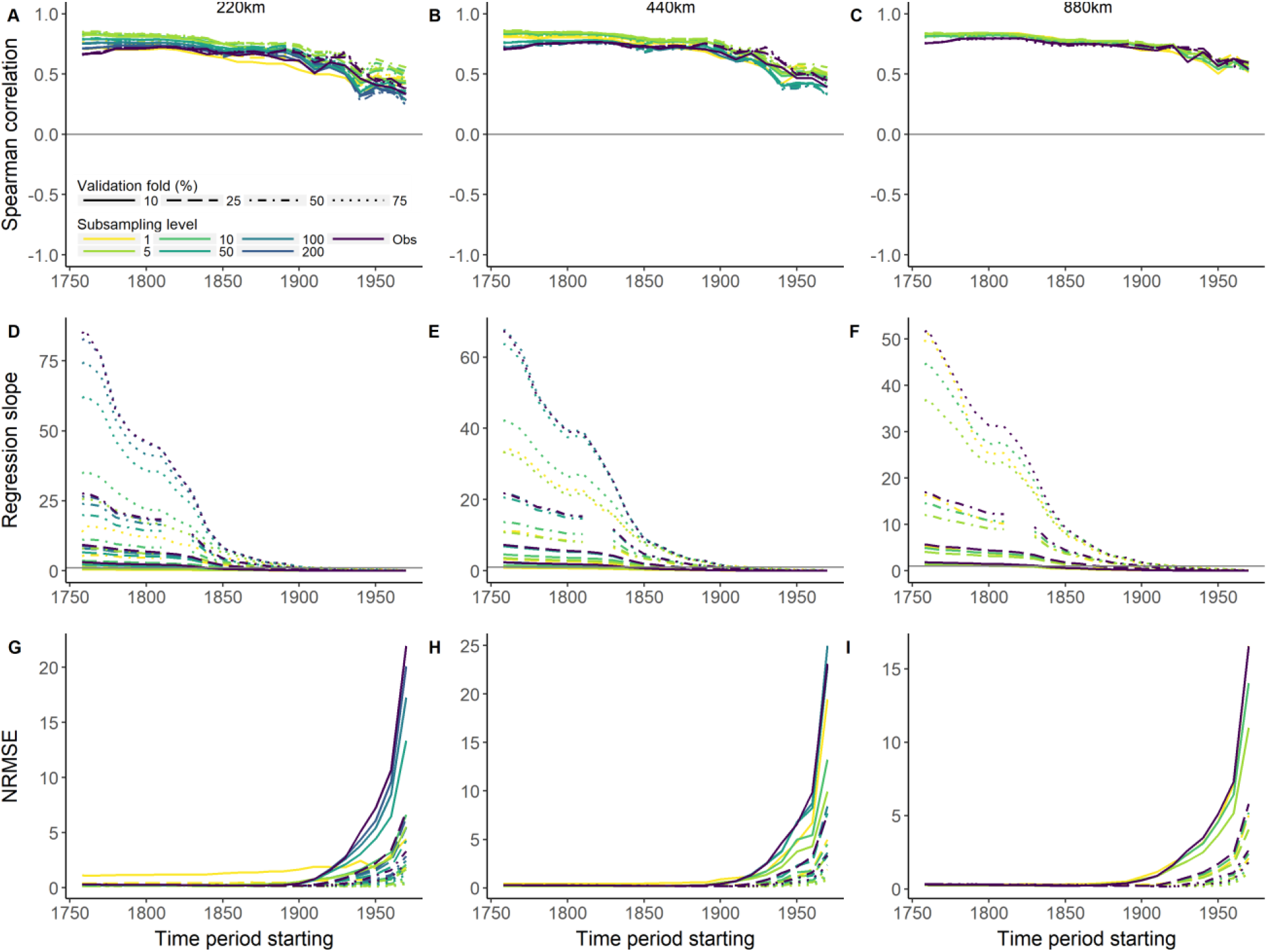
Statistic metrics derived from the sensitivity analysis using mammal species. All statistics were computed between estimated and observed species across grid cells. Line colours denote the subsampling level adopted to control overrepresentation of wide-ranging species. Panel columns refer to statistics calculated at different spatial resolutions (220, 440, 880 km). (A-C) Spearman correlation, (D-F) Regression slope, (G-I) Normalized Root Mean Square Error (NRMSE). Confidence intervals were omitted to increase readability.

**Fig. S12.**
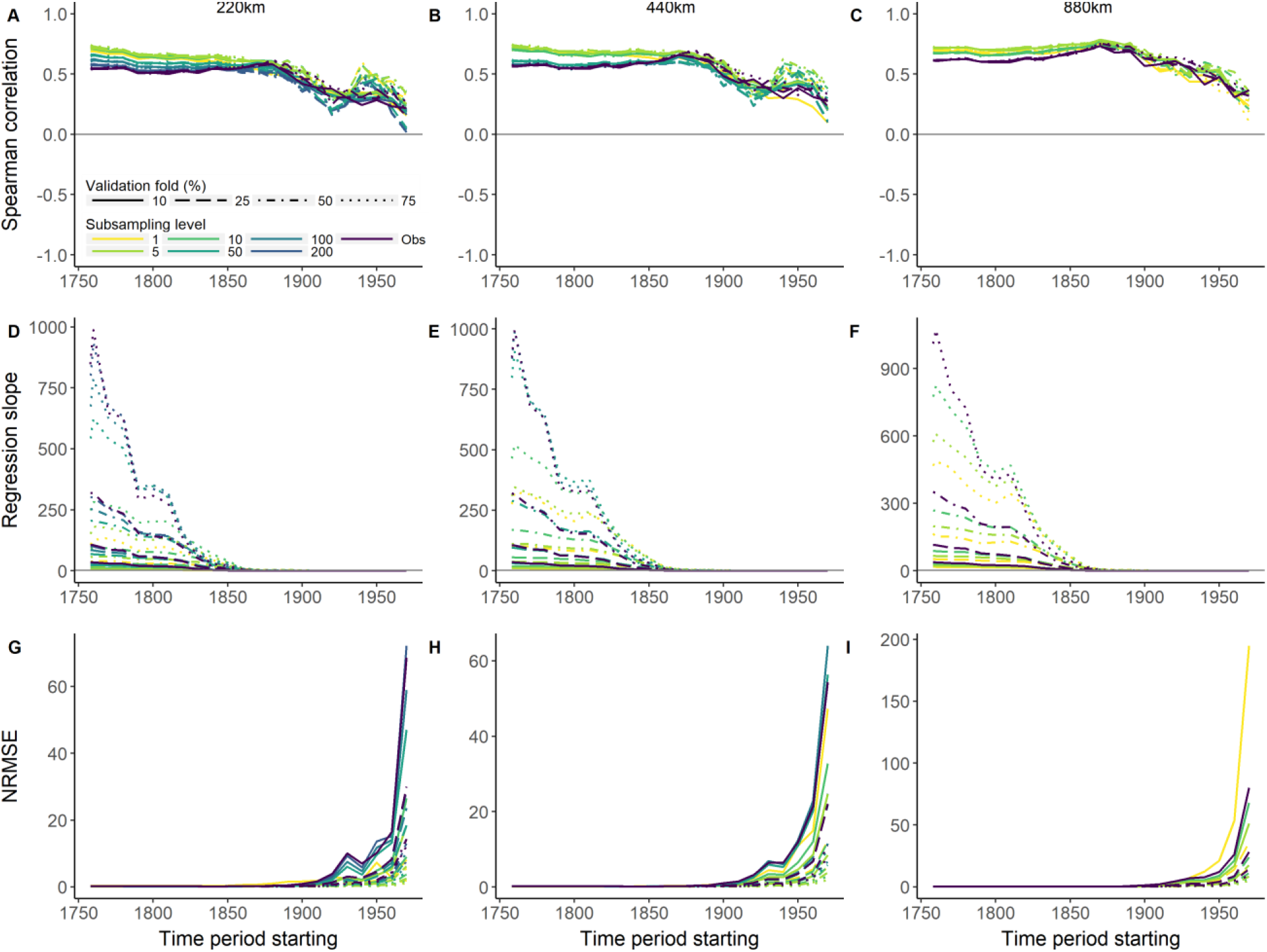
Statistic metrics derived from the sensitivity analysis using bird species. All statistics were computed between estimated and observed species across grid cells. Line colours denote the subsampling level adopted to control overrepresentation of wide-ranging species. Panel columns refer to statistics calculated at different spatial resolutions (220, 440, 880 km). (A-C) Spearman correlation, (D-F) Regression slope, (G-I) Normalized Root Mean Square Error (NRMSE). Confidence intervals were omitted to increase readability.

**Fig. S13.**
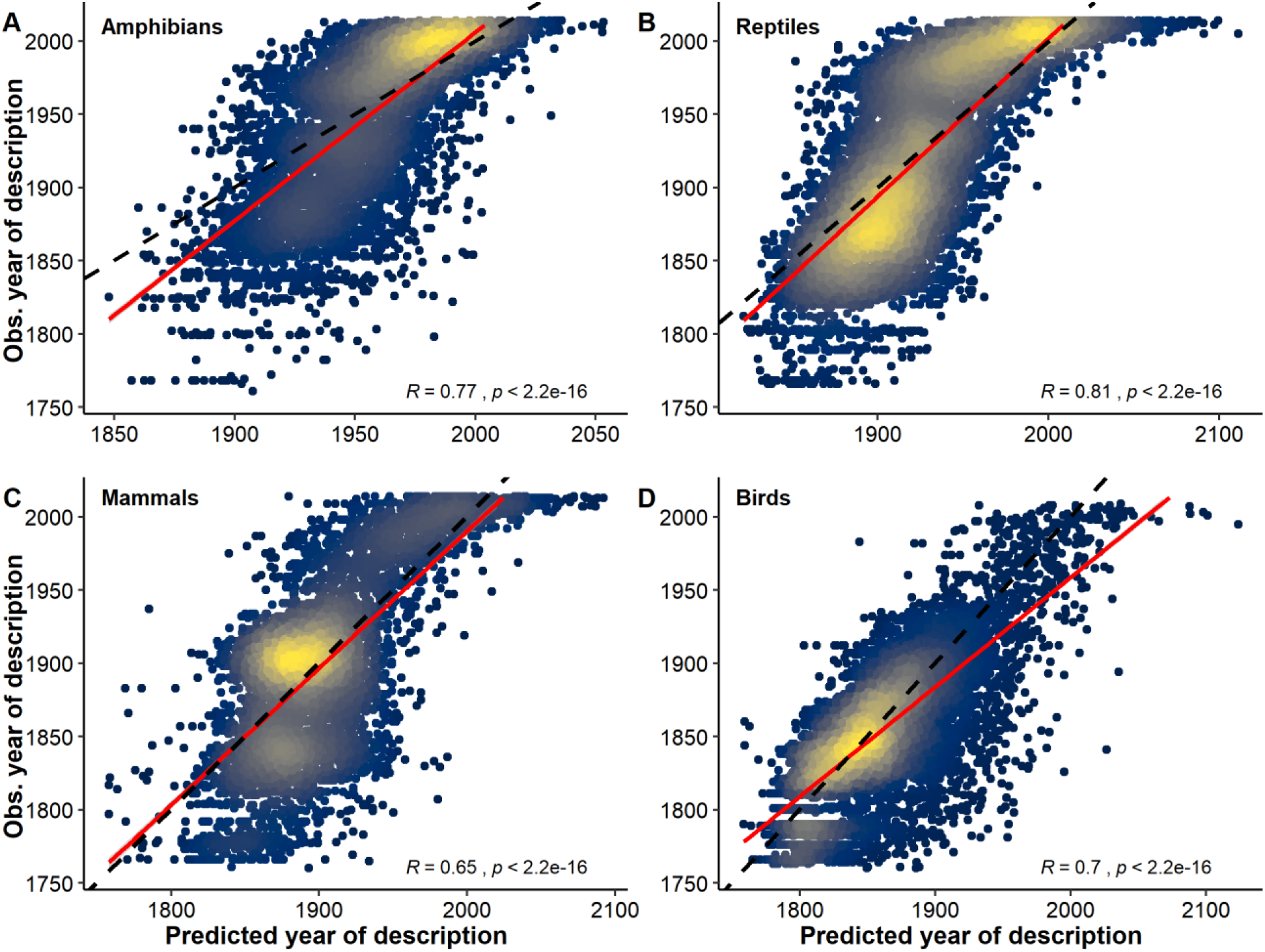
Relationship between predicted and observed year of description for terrestrial vertebrates. Plots include known species described from 1759 to 2014. (A) Amphibians, (B) Reptiles, (C) Mammals, (D) Birds. Models were first trained using 75% of the data, and then applied to the validation-fold (independent data) to obtain the predicted year of discovery for each species. The dashed line indicates the line of equality. R values inside plots denote the Spearman correlation between observed and predicted year of description.

**Fig. S14.**
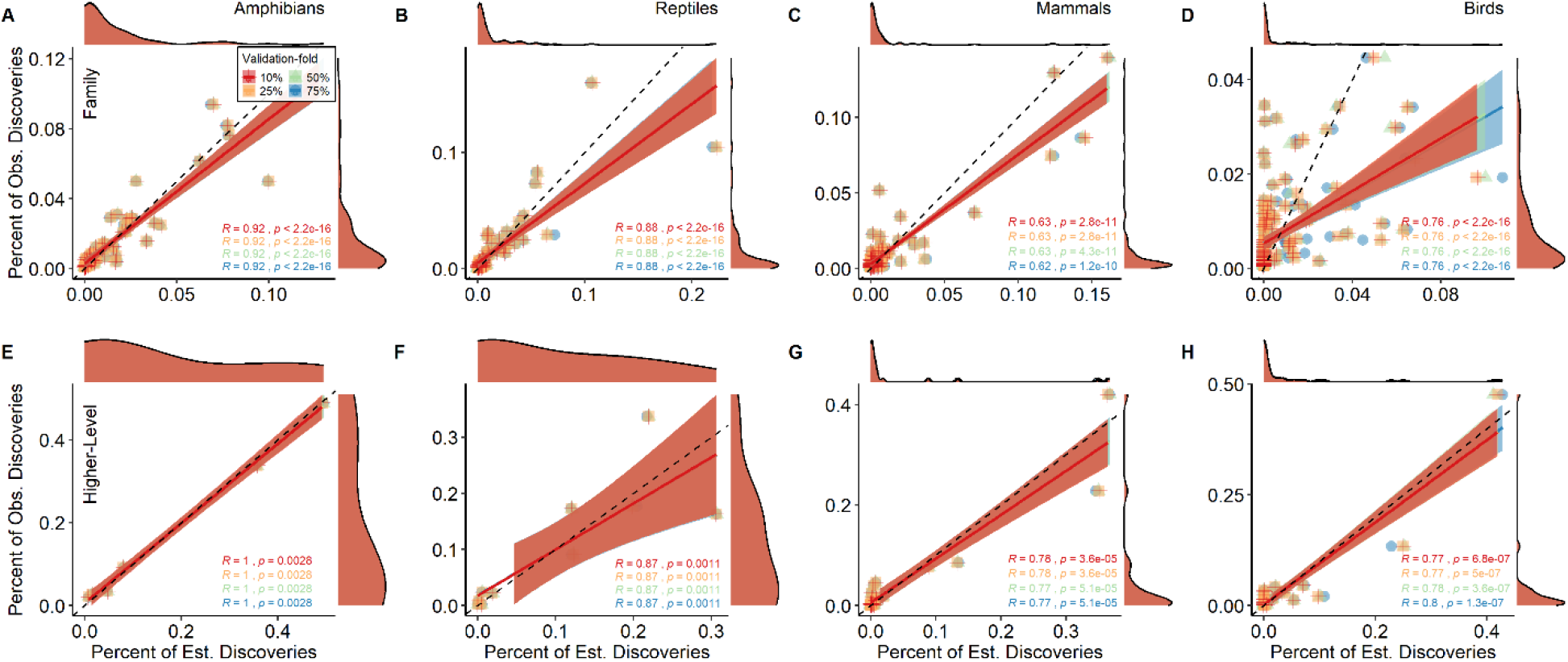
Relationship between estimated and observed discoveries at the taxon-level. Colours denote different sizes of the validation-fold. Columns represent different vertebrate groups and rows indicate different taxonomic ranks. The dashed line indicates the line of equality. (A, E, I) Amphibians. (B, F, J) Reptiles. (C, G, K) Mammals. (D, H, L) Birds. See ‘Model validation’ section for details on the highest-level taxonomic rank used. R values inside plots denote the Spearman correlation between estimated and observed discoveries. Plots include known species described from 1759 to 2014.

**Fig. S15.**
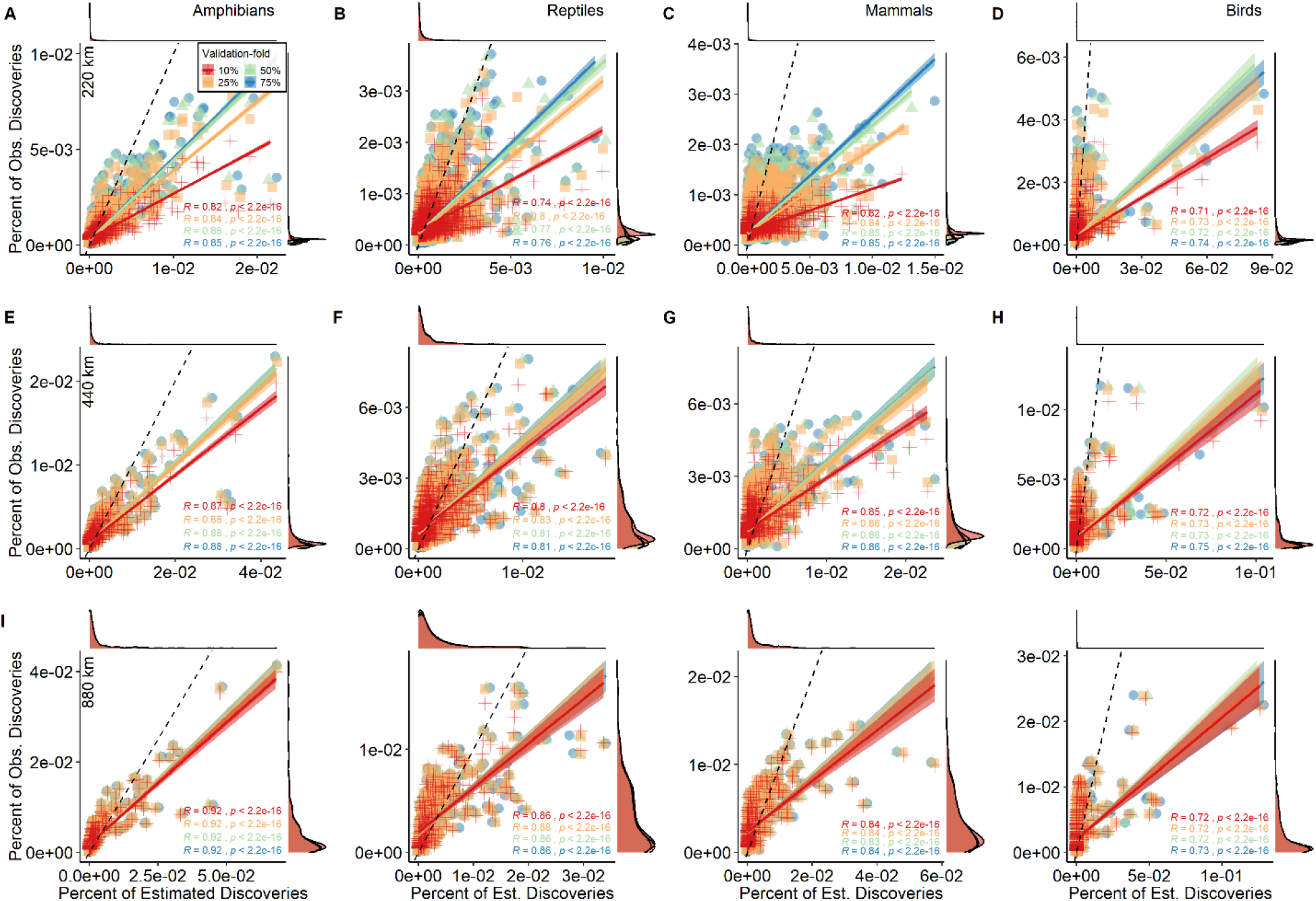
Relationship between estimated and observed discoveries at the assemblage-level. Colours denote different sizes of the validation-fold. Columns represent different vertebrate groups and rows indicate assemblages (grid cells) defined at different spatial resolutions. The dashed line indicates the line of equality. (A, E, I, M) Amphibians. (B, F, J, N) Reptiles. (C, G, K, O) Mammals. (D, H, L, P) Birds. Only the subsampling level of 5 is showed. R values inside plots denote the Spearman correlation between estimated and observed discoveries.

**Fig. S16.**
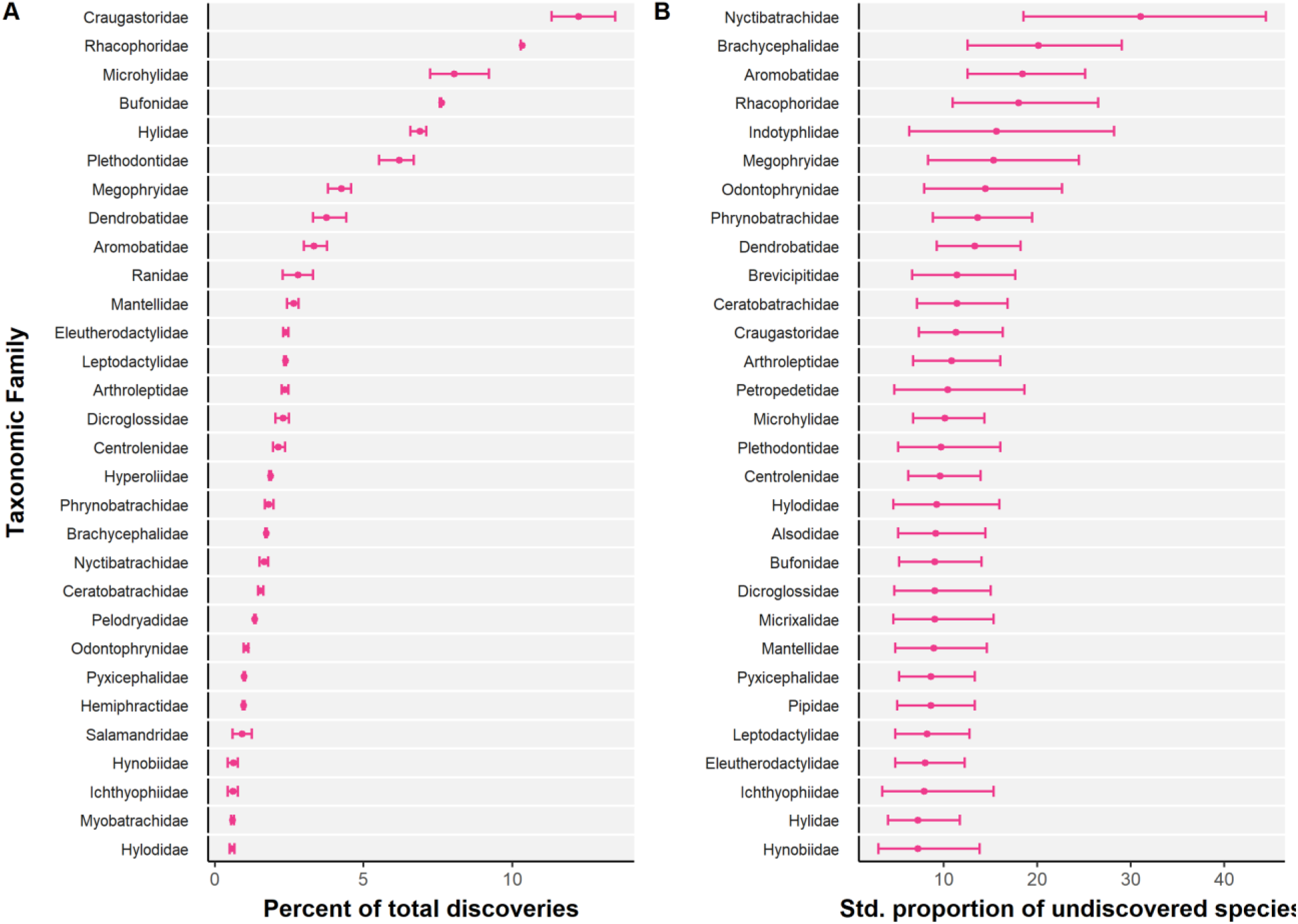
Top 30 amphibian families with highest potential for species discoveries. (A) Taxa with highest percent of total discoveries. (B) Taxa with highest standardized proportion of undiscovered species. The horizontal lines denote the 95% confidence intervals.

**Fig. S17.**
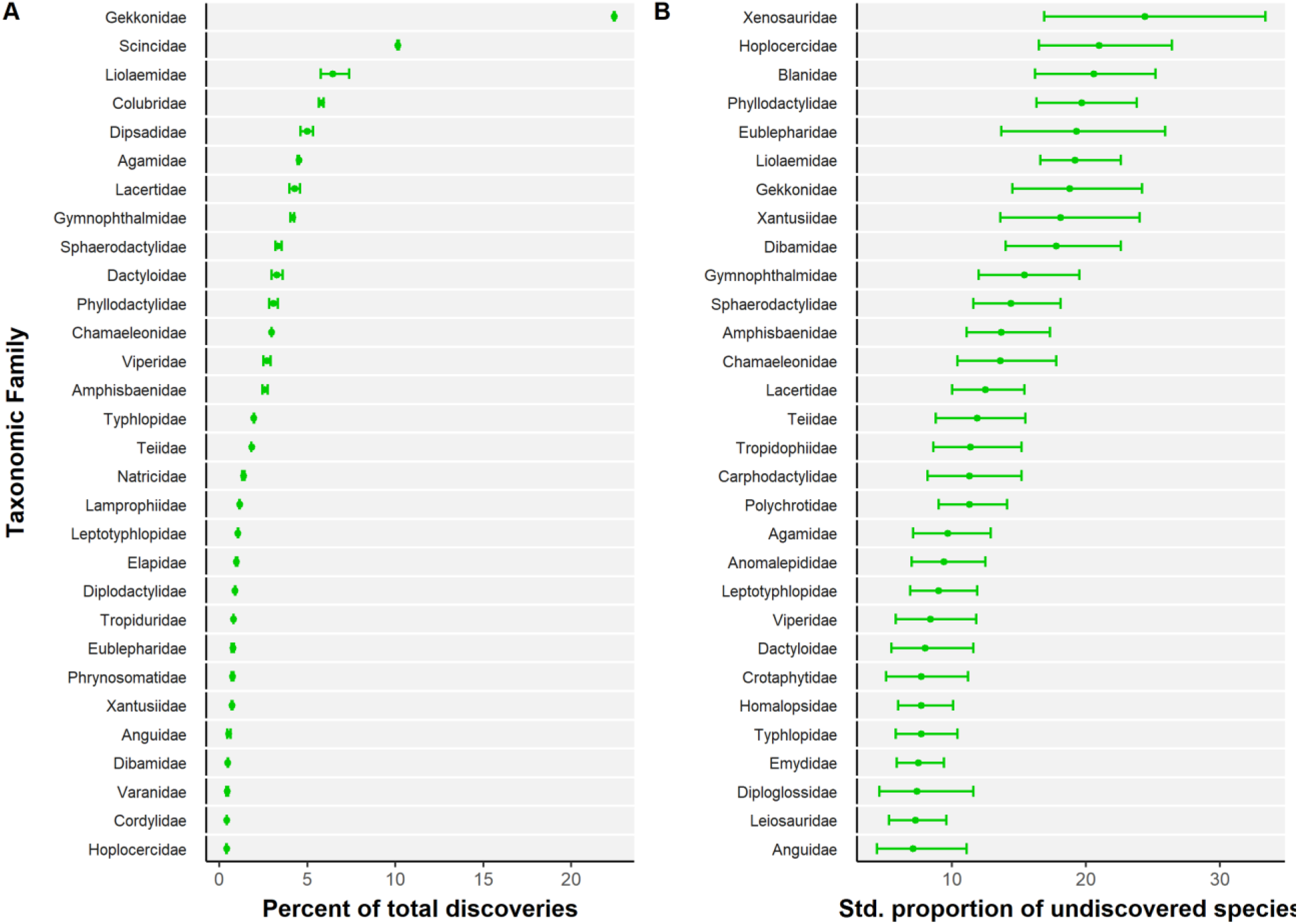
Top 30 reptile families with highest potential for species discoveries. (A) Taxa with highest percent of total discoveries. (B) Taxa with highest standardized proportion of undiscovered species. The horizontal lines denote the 95% confidence intervals.

**Fig. S18.**
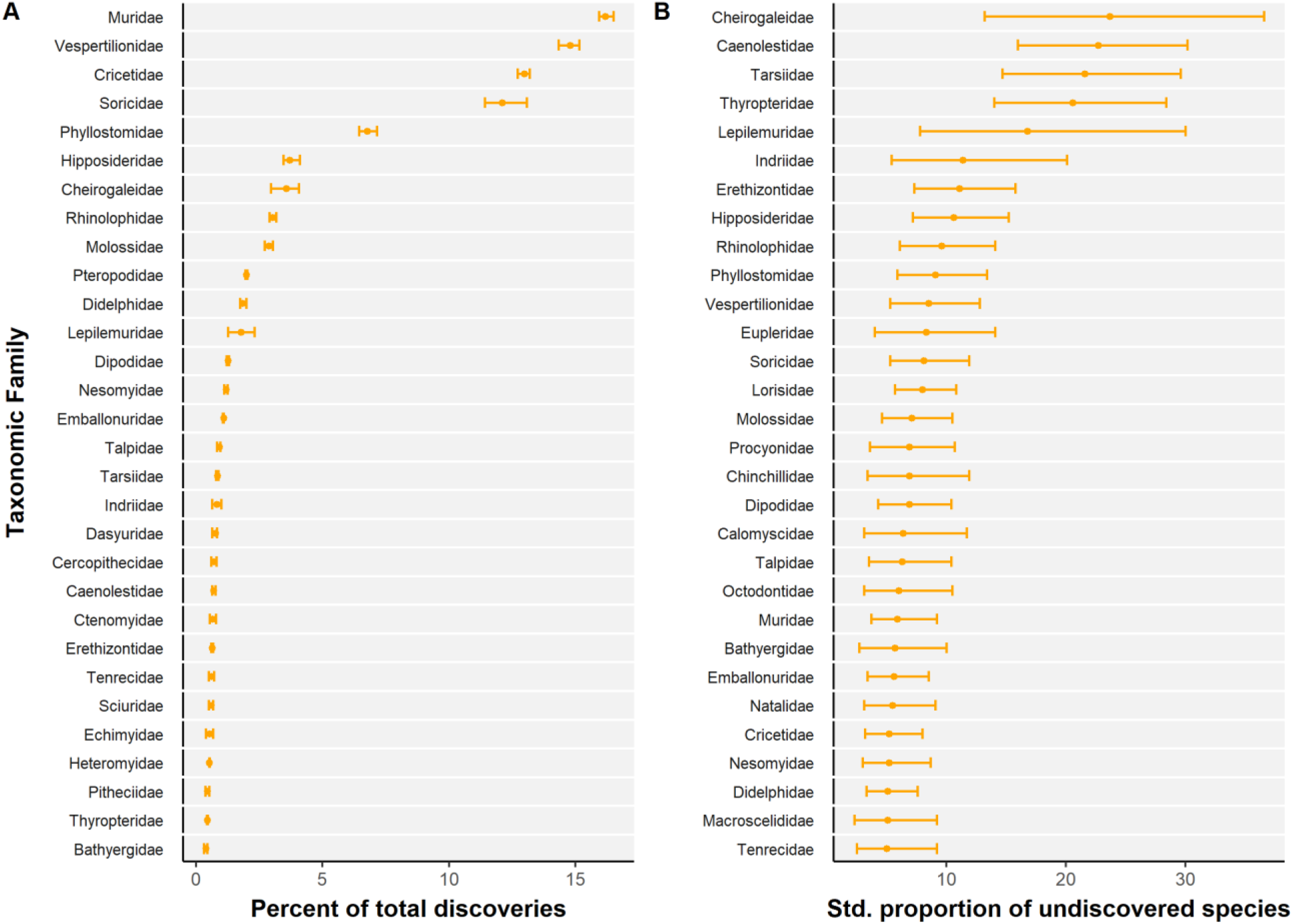
Top 30 mammal families with highest potential for species discoveries. (A) Taxa with highest percent of total discoveries. (B) Taxa with highest standardized proportion of undiscovered species. The horizontal lines denote the 95% confidence intervals.

**Fig. S19.**
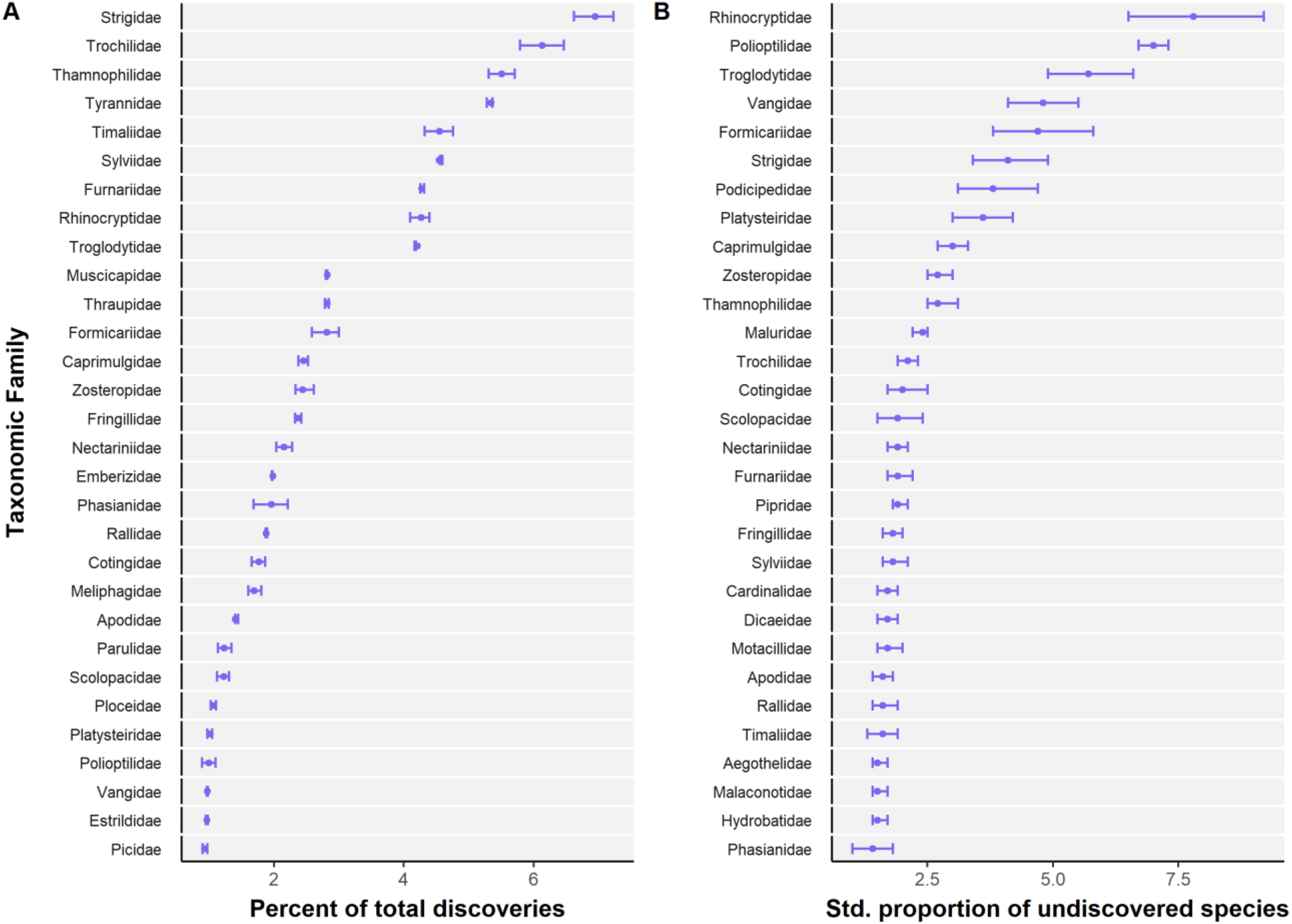
Top 30 bird families with highest potential for species discoveries. (A) Taxa with highest percent of total discoveries. (B) Taxa with highest standardized proportion of undiscovered species. The horizontal lines denote the 95% confidence intervals.

**Fig. S20.**
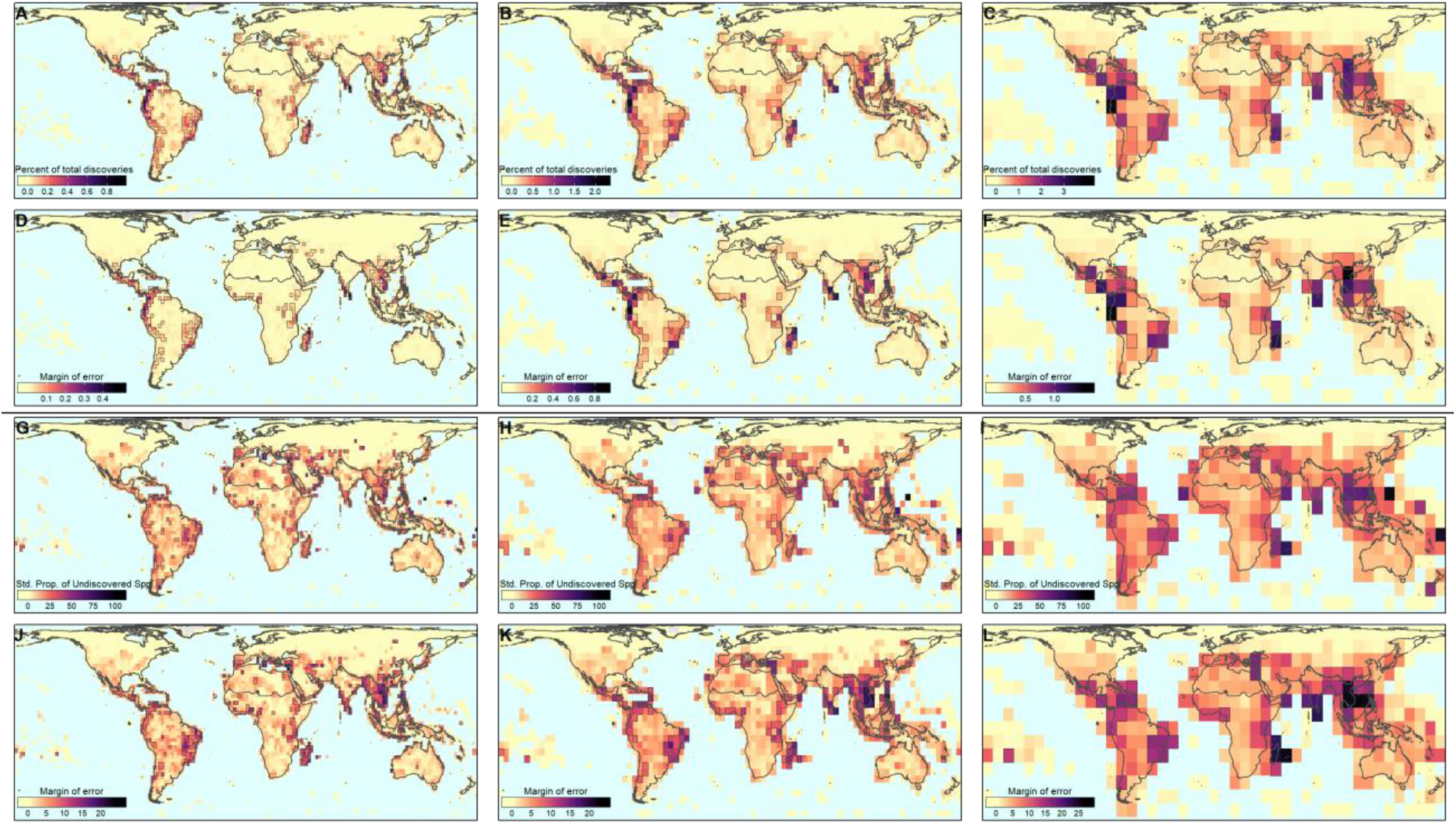
Geographical discovery patterns for terrestrial vertebrates at different spatial resolutions. (A-C) Percent of total predicted discoveries across grid cells and their respective (D-F) uncertainty (± margin of error). (G-I) Standardized proportion of undiscovered species across grid cells and their respective (J-L) uncertainty (± margin of error). Outlined and hatched regions designate grid cells holding values within respectively the top 10% and top 5% of the mapped metric. Maps drawn at spatial resolutions of 220, 440, 880 km.

**Fig. S21.**
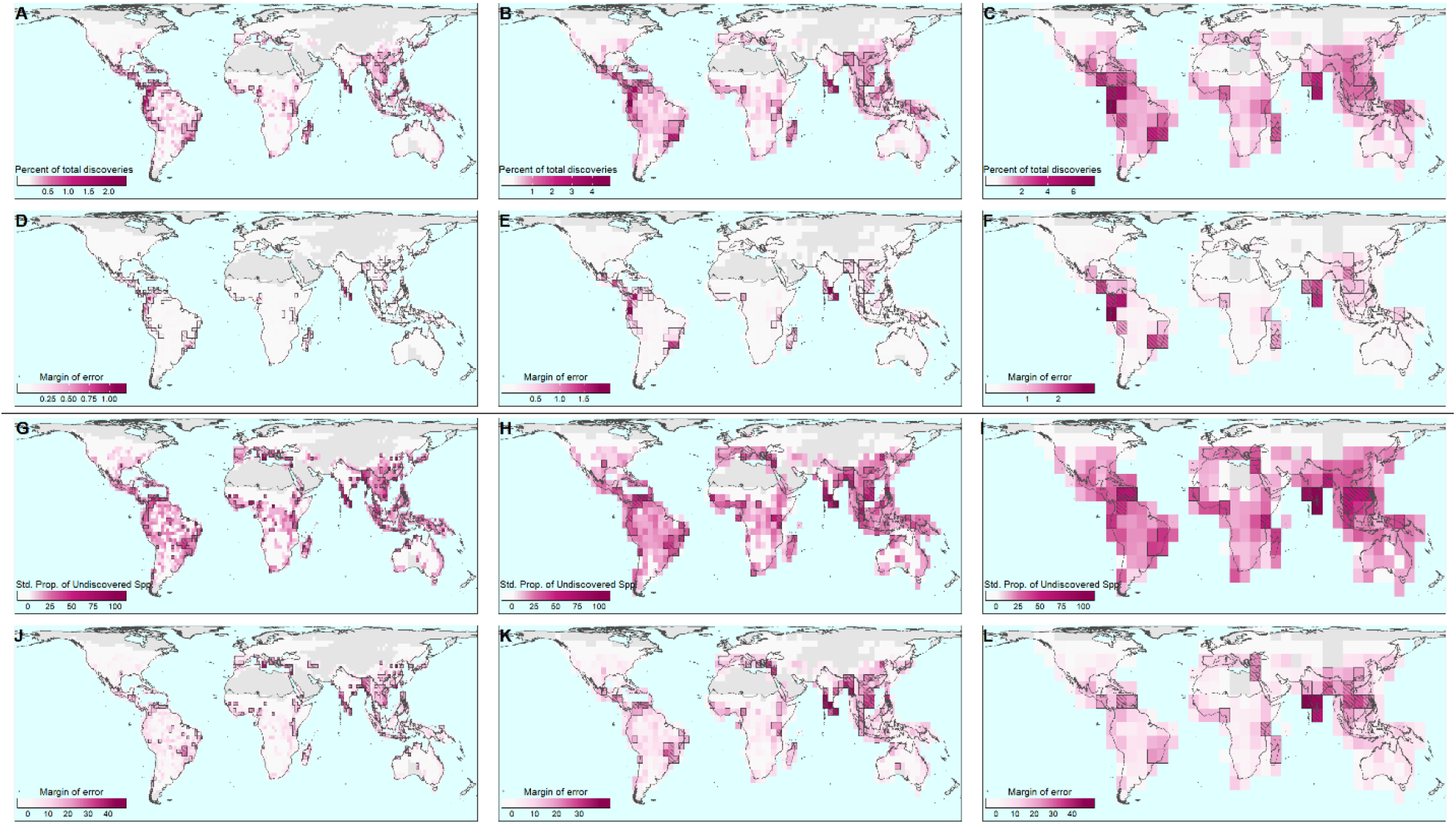
Geographical discovery patterns for amphibians at different spatial resolutions. (A-C) Percent of total discoveries across grid cells and their respective (D-F) uncertainty (± margin of error). (G-I) Standardized proportion of undiscovered species across grid cells and their respective (J-L) uncertainty (± margin of error). Outlined and hatched regions designate grid cells holding values within respectively the top 10% and top 5% of the mapped metric. Maps drawn at spatial resolutions of 220, 440, 880 km.

**Fig. S22.**
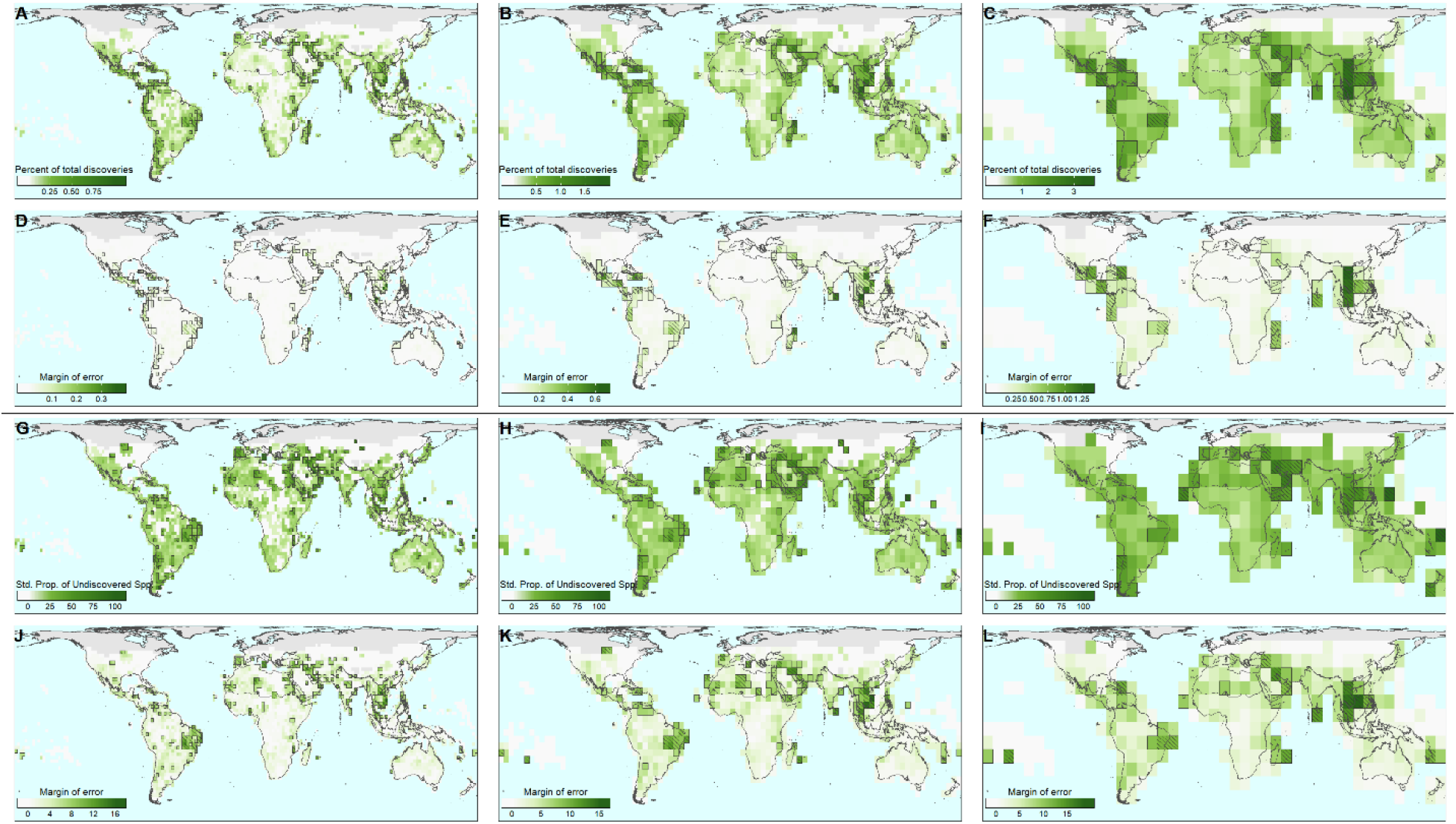
Geographical discovery patterns for reptiles at different spatial resolutions. (A-C) Percent of total discoveries across grid cells and their respective (D-F) uncertainty (± margin of error). (G-I) Standardized proportion of undiscovered species across grid cells and their respective (J-L) uncertainty (± margin of error). Outlined and hatched regions designate grid cells holding values within respectively the top 10% and top 5% of the mapped metric. Maps drawn at spatial resolutions of 220, 440, 880 km.

**Fig. S23.**
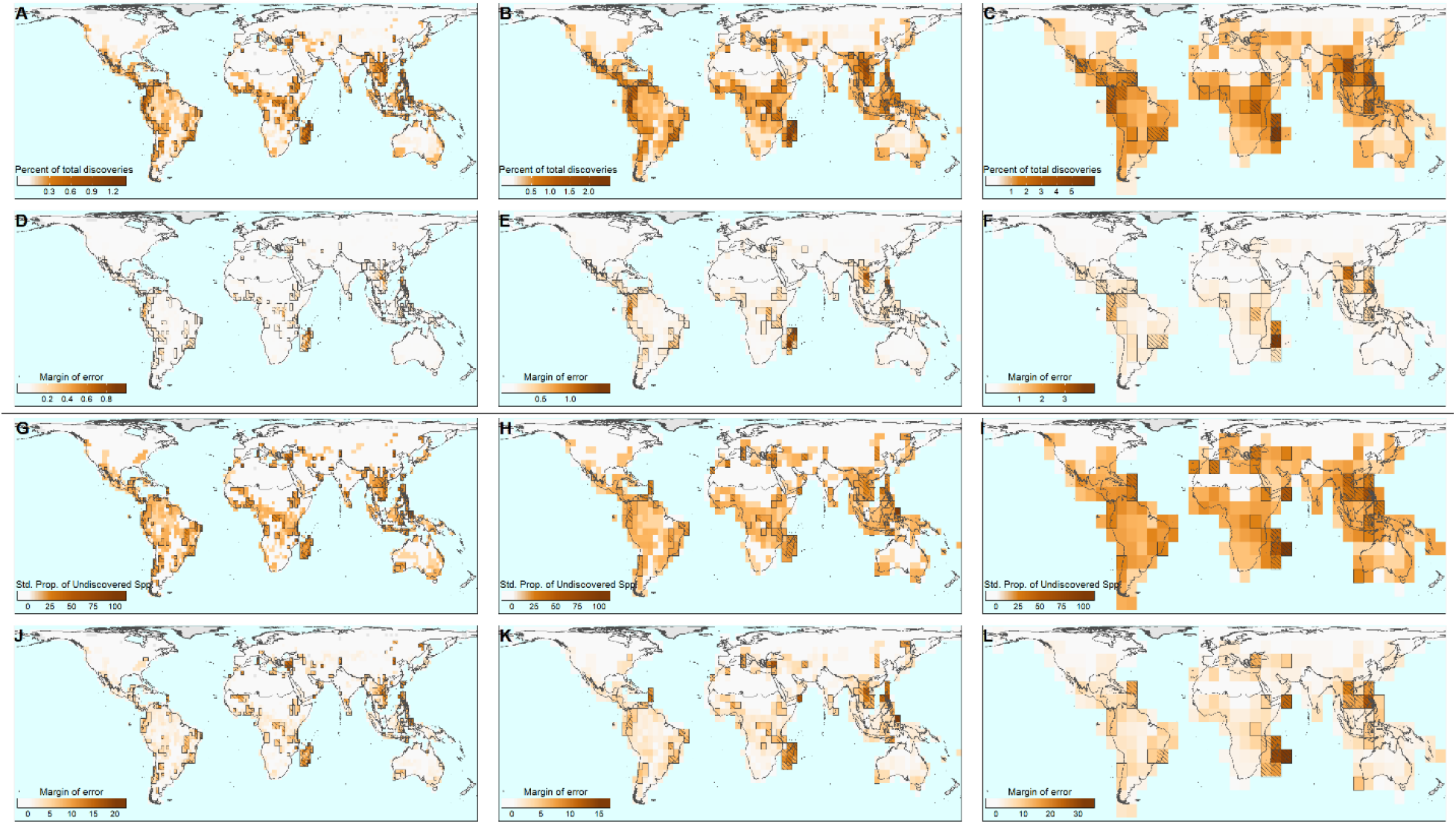
Geographical discovery patterns for mammals at different spatial resolutions. (A-C) Percent of total discoveries across grid cells and their respective (D-F) uncertainty (± margin of error). (G-I) Standardized proportion of undiscovered species across grid cells and their respective (J-L) uncertainty (± margin of error). Outlined and hatched regions designate grid cells holding values within respectively the top 10% and top 5% of the mapped metric. Maps drawn at spatial resolutions of 220, 440, 880 km.

**Fig. S24.**
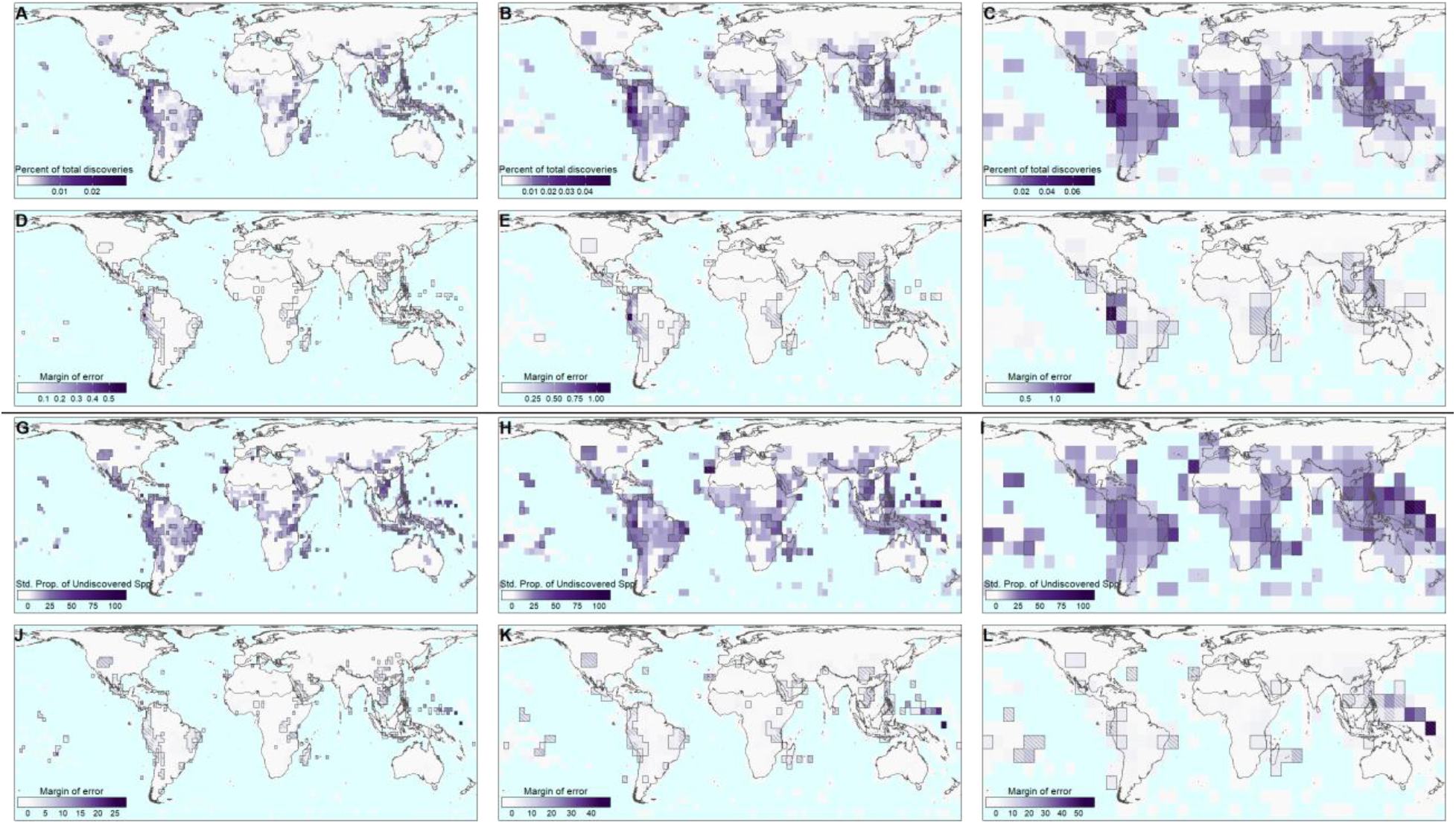
Geographical discovery patterns for birds at different spatial resolutions. (A-C) Percent of total discoveries across grid cells and their respective (D-F) uncertainty (± margin of error). (G-I) Standardized proportion of undiscovered species across grid cells and their respective (J-L) uncertainty (± margin of error). Outlined and hatched regions designate grid cells holding values within respectively the top 10% and top 5% of the mapped metric. Maps drawn at spatial resolutions of 220, 440, 880 km.

**Fig. S25.**
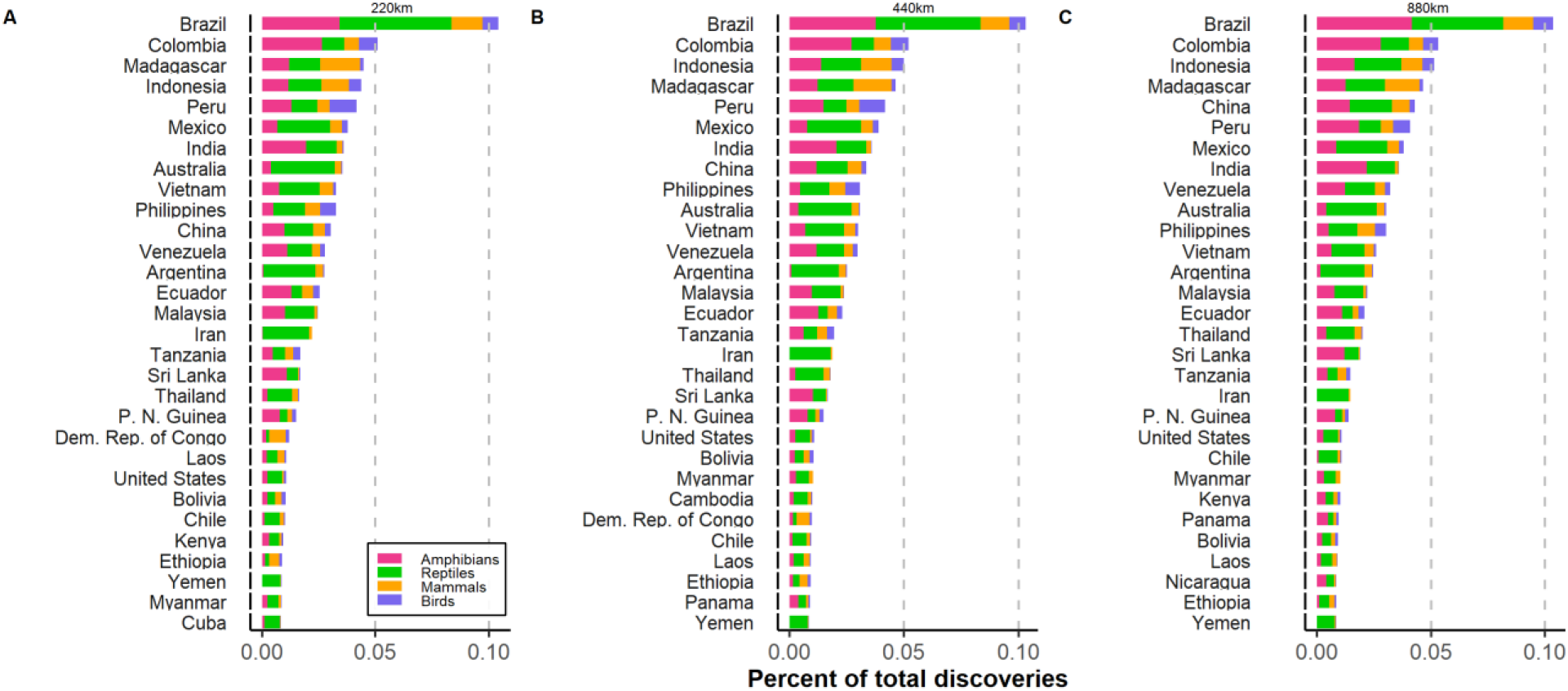
Top 30 countries with higher percent of total discoveries. Country-wide percent of total discoveries extracted from assemblages defined at (A) 220 km, (B) 440 km, and (C) 880 km of spatial resolution.

## REFERENCES

1. Costello, M. J., May, R. M. & Stork, N. E. Can We Name Earth’s Species Before They Go Extinct? Science 339, 413–416 (2013).

2. Mora, C., Rollo, A. & Tittensor, D. P. Comment on ‘Can We Name Earth’s Species Before They Go Extinct?’ Science 341, 237 (2013).

3. Mora, C., Tittensor, D. P., Adl, S., Simpson, A. G. B. & Worm, B. How Many Species Are There on Earth and in the Ocean? PLoS Biol. 9, e1001127 (2011).

4. May, R. & Beverton, R. J. H. How many species? Philos. Trans. R. Soc. London. Ser. B Biol. Sci. 330, 293–304 (1990).

5. Scheffers, B. R., Joppa, L. N., Pimm, S. L. & Laurance, W. F. What we know and don’t know about Earth’s missing biodiversity. Trends Ecol. Evol. 27, 501–510 (2012).

6. Raven, P. H. & Wilson, E. O. A fifty-year plan for biodiversity surveys. Science 258, 1099–1100 (1992).

7. Whittaker, R. J. et al. Conservation biogeography: Assessment and prospect. Divers. Distrib. 11, 3–23 (2005).

8. Hortal, J. et al. Seven Shortfalls that Beset Large-Scale Knowledge of Biodiversity. Annu. Rev. Ecol. Evol. Syst. 46, 523–549 (2015).

9. Secretariat of the Convention on Biological Diversity. Guide to the global taxonomy initiative. CBD Tech. Ser. 30, 1–195, i–viii (2010).

10. Costello, M. J., May, R. M. & Stork, N. E. Response to Comments on ‘Can We Name Earth’s Species Before They Go Extinct?’ Science 341, 237 (2013).

11. Bebber, D. P., Marriott, F. H. C., Gaston, K. J., Harris, S. A. & Scotland, R. W. Predicting unknown species numbers using discovery curves. Proc. R. Soc. B 274, 1651–1658 (2007).

12. Edie, S. M., Smits, P. D. & Jablonski, D. Probabilistic models of species discovery and biodiversity comparisons. Proc. Natl. Acad. Sci. 114, 3666–3671 (2017).

13. Guenard, B., Weiser, M. D. & Dunn, R. R. Global models of ant diversity suggest regions where new discoveries are most likely are under disproportionate deforestation threat. Proc. Natl. Acad. Sci. 109, 7368–7373 (2012).

14. Blackburn, T. M. & Gaston, K. J. What Determines the Probability of Discovering a Species - a Study of South-American Oscine Passerine Birds. J. Biogeogr. 22, 7–14 (1995).

15. Costello, M. J., Lane, M., Wilson, S. & Houlding, B. Factors influencing when species are first named and estimating global species richness. Glob. Ecol. Conserv. 4, 243–254 (2015).

16. Collen, B., Purvis, A. & Gittleman, J. L. Biological correlates of description date in carnivores and primates. Glob. Ecol. Biogeogr. 13, 459–467 (2004).

17. Diniz-Filho, J. A. F. et al. Macroecological correlates and spatial patterns of anuran description dates in the Brazilian Cerrado. Glob. Ecol. Biogeogr. 14, 469–477 (2005).

18. Costello, M. J., Houlding, B. & Joppa, L. N. Further evidence of more taxonomists discovering new species, and that most species have been named: response to Bebber et al. (2014). New Phytol. 202, 739–740 (2014).

19. Meiri, S. Small, rare and trendy: traits and biogeography of lizards described in the 21st century. J. Zool. 299, 251–261 (2016).

20. Diniz-Filho, J. A. F. et al. Macroecological correlates and spatial patterns of anuran description dates in the Brazilian Cerrado. Glob. Ecol. Biogeogr. 14, 469–477 (2005).

21. Klein, J. P. & Moeschberger, M. L. Survival analysis: Techniques for censored and truncated data. Pharmaceutical Statistics (Springer, 2003). doi:10.1002/pst.135

22. Essl, F., Rabitsch, W., Dullinger, S., Moser, D. & Milasowszky, N. How well do we know species richness in a well-known continent? Temporal patterns of endemic and widespread species descriptions in the European fauna. Glob. Ecol. Biogeogr. 22, 29–39 (2013).

23. Colli, G. R. et al. In the depths of obscurity: Knowledge gaps and extinction risk of Brazilian worm lizards (Squamata, Amphisbaenidae). Biol. Conserv. 204, 51–62 (2016).

24. Burgin, C. J., Colella, J. P., Kahn, P. L. & Upham, N. S. How many species of mammals are there? J. Mammal. 99, 1–14 (2018).

25. Meyer, C., Kreft, H., Guralnick, R. & Jetz, W. Global priorities for an effective information basis of biodiversity distributions. Nat. Commun. 6, 8221 (2015).

26. Bellard, C. et al. Vulnerability of biodiversity hotspots to global change. Glob. Ecol. Biogeogr. 23, 1376–1386 (2014).

27. Quintero, I. & Jetz, W. Global elevational diversity and diversification of birds. Nature 555, 246–250 (2018).

28. Roll, U. et al. The global distribution of tetrapods reveals a need for targeted reptile conservation. Nat. Ecol. Evol. 1, 1677–1682 (2017).

29. Garnett, S. T. & Christidis, L. Taxonomy anarchy hampers conservation. Nature 546, 25–27 (2017).

30. Isaac, N. J. B., Mallet, J. & Mace, G. M. Taxonomic inflation: its influence on macroecology and conservation. Trends Ecol. Evol. 19, 464–469 (2004).

31. Bremer, K., Bremer, B., Karis, P. & Källersjö, M. Time for change in taxonomy. Nature 343, 202–202 (1990).

32. Raposo, M. A. et al. What really hampers taxonomy and conservation? A riposte to Garnett and Christidis (2017). Zootaxa 4317, 179 (2017).

33. Wake, D. B. Persistent Plethodontid Themes: Species, Phylogenies, and Biogeography. Herpetologica 73, 242–251 (2017).

34. Tedesco, P. A. et al. Estimating How Many Undescribed Species Have Gone Extinct. Conserv. Biol. 28, 1360–1370 (2014).

35. Jetz, W., McPherson, J. M. & Guralnick, R. P. Integrating biodiversity distribution knowledge: toward a global map of life. Trends Ecol. Evol. 27, 151–159 (2012).

36. Jetz, W., Thomas, G. H., Joy, J. B., Hartmann, K. & Mooers, A. O. The global diversity of birds in space and time. Nature 491, 444–448 (2012).

37. Jetz, W. & Pyron, R. A. The interplay of past diversification and evolutionary isolation with present imperilment across the amphibian tree of life. Nat. Ecol. Evol. 2, 850–858 (2018).

38. Upham, N. S., Esselstyn, J. A. & Jetz, W. Inferring the mammal tree: Species-level sets of phylogenies for questions in ecology, evolution, and conservation. PLOS Biol. 17, e3000494 (2019).

39. González-del-Pliego, P. et al. Phylogenetic and Trait-Based Prediction of Extinction Risk for Data-Deficient Amphibians. Curr. Biol. 29, 1557–1563.e3 (2019).

40. Moura, M. R. et al. Geographical and socioeconomic determinants of species discovery trends in a biodiversity hotspot. Biol. Conserv. 220, 237–244 (2018).

41. Gaston, K. J., Blackburn, T. M. & Loder, N. Which species are described first? The case of North-American butterflies. Biodivers. Conserv. 4, 119–127 (1995).

42. Oliveira, B. F., São-Pedro, V. A., Santos-Barrera, G., Penone, C. & Costa, G. C. AmphiBIO, a global database for amphibian ecological traits. Sci. Data 4, 170123 (2017).

43. Feldman, A., Sabath, N., Pyron, R. A., Mayrose, I. & Meiri, S. Body sizes and diversification rates of lizards, snakes, amphisbaenians and the tuatara. Glob. Ecol. Biogeogr. 25, 187–197 (2016).

44. Hallmann, K. & Griebeler, E. M. An exploration of differences in the scaling of life history traits with body mass within reptiles and between amniotes. Ecol. Evol. 8, 5480–5494 (2018).

45. Slavenko, A., Itescu, Y., Ihlow, F. & Meiri, S. Home is where the shell is: predicting turtle home range sizes. J. Anim. Ecol. 85, 106–114 (2016).

46. Regis, K. W. & Meik, J. M. Allometry of sexual size dimorphism in turtles: a comparison of mass and length data. PeerJ 5, e2914 (2017).

47. Itescu, Y., Karraker, N. E., Raia, P., Pritchard, P. C. H. & Meiri, S. Is the island rule general? Turtles disagree. Glob. Ecol. Biogeogr. 23, 689–700 (2014).

48. Upham, N. S., Esselstyn, J. A. & Jetz, W. Ecological causes of uneven diversification and richness in the mammal tree of life. bioRxiv (2019). doi:10.1101/504803

49. Wilman, H. et al. EltonTraits 1.0: Species-level foraging attributes of the world’s birds and mammals. Ecology 95, 2027–2027 (2014).

50. Tonini, J. F. R., Beard, K. H., Ferreira, R. B., Jetz, W. & Pyron, R. A. Fully-sampled phylogenies of squamates reveal evolutionary patterns in threat status. Biol. Conserv. (2016). doi:10.1016/j.biocon.2016.03.039

51. Goolsby, E. W., Bruggeman, J. & Ané, C. Rphylopars : fast multivariate phylogenetic comparative methods for missing data and within-species variation. Methods Ecol. Evol. 8, 22–27 (2017).

52. Gaston, K. J., Blackburn, T. M. & Lawton, J. H. Interspecific Abundance-Range Size Relationships: An Appraisal of Mechanisms. J. Anim. Ecol. 66, 579 (1997).

53. Borregaard, M. K. & Rahbek, C. Causality of the Relationship between Geographic Distribution and Species Abundance. Q. Rev. Biol. 85, 3–25 (2010).

54. IUCN - International Union for Conservation of Nature. IUCN Red List of Threatened Species. Version 2018 www.iucnredlist.org (2018).

55. Freitag, S., Hobson, C., Biggs, H. C. & Jaarsveld, A. S. Testing for potential survey bias: the effect of roads, urban areas and nature reserves on a southern African mammal data set. Anim. Conserv. 1, 119–127 (1998).

56. Kier, G. & Barthlott, W. Measuring and mapping endemism and species richness: a new methodological approach and its application on the flora of Africa. Biodivers. Conserv. 10, 1513–1529 (2001).

57. Vilela, B. & Villalobos, F. letsR: a new R package for data handling and analysis in macroecology. Methods Ecol. Evol. 6, 1229–1234 (2015).

58. Papavero, N. Essays on the History of Neotropical Dipterology: with special reference to collectors: 1750–1905: Vol. I. (Museu de Zoologia da Universidade de São Paulo, 1971). doi:10.5962/bhl.title.101715

59. Baselga, A., Lobo, J. M., Hortal, J., Jiménez-Valverde, A. & Gómez, J. F. Assessing alpha and beta taxonomy in eupelmid wasps: determinants of the probability of describing good species and synonyms. J. Zool. Syst. Evol. Res. 48, 40–49 (2010).

60. Yang, W., Ma, K. & Kreft, H. Environmental and socio-economic factors shaping the geography of floristic collections in China. Glob. Ecol. Biogeogr. 23, 1284–1292 (2014).

61. Karger, D. N. et al. Climatologies at high resolution for the earth’s land surface areas. Sci. Data 4, 170122 (2017).

62. R Core Team. R: A Language and Environment for Statistical Computing. v. 3.5.3 (2019).

63. Hijmans, R. J. raster: Geographic Data Analysis and Modeling. https://cran.r-project.org/package=raster (2015).

64. Amatulli, G. et al. A suite of global, cross-scale topographic variables for environmental and biodiversity modeling. Sci. Data 5, 180040 (2018).

65. Klein Goldewijk, K., Beusen, A., Van Drecht, G. & De Vos, M. The HYDE 3.1 spatially explicit database of human-induced global land-use change over the past 12,000 years. Glob. Ecol. Biogeogr. 20, 73–86 (2011).

66. Joppa, L. N., Roberts, D. L. & Pimm, S. L. The population ecology and social behaviour of taxonomists. Trends Ecol. Evol. 26, 551–553 (2011).

67. Wickham, H. stringr: Simple, Consistent Wrappers for Common String Operations. R package version 1.3.1. http://stringr.tidyverse.org (2018).

68. Mahto, A. splitstackshape: Stack and Reshape Datasets After Splitting Concatenated Values. R package version 1.4.6. http://github.com/mrdwab/splitstackshape (2018).

69. Dinerstein, E. et al. An Ecoregion-Based Approach to Protecting Half the Terrestrial Realm. Bioscience 67, 534–545 (2017).

70. Jetz, W. & Fine, P. V. A. Global Gradients in Vertebrate Diversity Predicted by Historical Area-Productivity Dynamics and Contemporary Environment. PLoS Biol. 10, e1001292 (2012).

71. Kutner, M. H., Nachtsheim, C. J., Neter, J. & Li, W. Applied Linear Statistical Models. (McGraw-Hill Irwin, 2004).

72. Naimi, B. usdm: Uncertainty Analysis for Species Distribution Models. https://cran.r-project.org/package=usdm (2017).

73. Bebber, D. P. et al. Herbaria are a major frontier for species discovery. Proc. Natl. Acad. Sci. 107, 22169–22171 (2010).

74. Guedes, J. J. M., Feio, R. N., Meiri, S. & Moura, M. R. Identifying factors that boost species discoveries of global reptiles. Zool. J. Linn. Soc. in press (2020). doi:10.1093/zoolinnean/zlaa029

75. von Linné, C. Caroli Linnaei…Systema naturae per regna tria naturae: secundum classes, ordines, genera, species, cum characteribus, differentiis, synonymis, locis. (Impensis Direct. Laurentii Salvii, 1758). doi:10.5962/bhl.title.542

76. Harrell, F. E. Regression Modeling Strategies. (Springer New York, 2001). doi:10.1007/978-1-4757-3462-1

77. George, B., Seals, S. & Aban, I. Survival analysis and regression models. J. Nucl. Cardiol. 21, 686–694 (2014).

78. Jackson, C. flexsurv : A Platform for Parametric Survival Modeling in R. J. Stat. Softw. 70, 1–33 (2016).

79. Burnham, K. P. & Anderson, D. R. Model Selection and Multimodel Inference. (Springer New York, 2004). doi:10.1007/b97636

80. Burnham, K. P. & Anderson, D. R. Model Selection and Multimodel Inference: A Practical Information-Theoretic Approach. Ecological Modelling 172, (Springer, 2002).

81. Johnson, J. B. & Omland, K. S. Model selection in ecology and evolution. Trends Ecol. Evol. 19, 101–108 (2004).

82. Barton, K. MuMIn: Multi-Model Inference. R package version 1.43.6. 1–74 (2019). Available at: https://cran.r-project.org/package=MuMIn.

83. Alexander Pyron, R. & Wiens, J. J. A large-scale phylogeny of Amphibia including over 2800 species, and a revised classification of extant frogs, salamanders, and caecilians. Mol. Phylogenet. Evol. 61, 543–583 (2011).

84. Pyron, R. A., Burbrink, F. T. & Wiens, J. J. A phylogeny and revised classification of Squamata, including 4161 species of lizards and snakes. BMC Evol. Biol. 13, 93 (2013).

85. Fisher, D. O. & Blomberg, S. P. Correlates of rediscovery and the detectability of extinction in mammals. Proc. R. Soc. B Biol. Sci. 278, 1090–1097 (2011).

86. Jetz, W., Sekercioglu, C. H. & Böhning-Gaese, K. The Worldwide Variation in Avian Clutch Size across Species and Space. PLoS Biol. 6, e303 (2008).

87. Jetz, W. & Rubenstein, D. R. Environmental Uncertainty and the Global Biogeography of Cooperative Breeding in Birds. Curr. Biol. 21, 72–78 (2011).

88. Jetz, W. & Rahbek, C. Geographic range size and determinants of avian species richness. Science 297, 1548–1551 (2002).

89. Dowle, M. & Srinivasan, A. data.table: Extension of ‘data.frame’. R package version 1.12.4. (2019). Available at: https://cran.r-project.org/package=data.table.

90. Gaston, K. J., Chown, S. L. & Evans, K. L. Ecogeographical rules: elements of a synthesis. J. Biogeogr. 35, 483–500 (2008).

91. Violle, C., Reich, P. B., Pacala, S. W., Enquist, B. J. & Kattge, J. The emergence and promise of functional biogeography. Proc. Natl. Acad. Sci. 111, 13690–13696 (2014).

92. GDAM. Database of Global Administrative Areas, version 3.6. (2019). Available at: http://www.gadm.org.

## SUPPLEMENTARY REFERENCES

1. Barnston, A. G. Correspondence among the Correlation, RMSE, and Heidke Forecast Verification Measures; Refinement of the Heidke Score. Weather Forecast. 7, 699–709 (1992).

2. Zambrano-Bigiarini, M. hydroGOF: Goodness-of-fit functions for comparison of simulated and observed hydrological time series. (2017). doi:10.5281/zenodo.840087

3. R Core Team. R: A Language and Environment for Statistical Computing. v. 3.5.3 (2019).

4. Bebber, D. P., Marriott, F. H. C., Gaston, K. J., Harris, S. A. & Scotland, R. W. Predicting unknown species numbers using discovery curves. Proc. R. Soc. B 274, 1651–1658 (2007).

5. Klein, J. P. & Moeschberger, M. L. Survival analysis: Techniques for censored and truncated data. Pharmaceutical Statistics (Springer, 2003). doi:10.1002/pst.135

6. Pante, E., Schoelinck, C. & Puillandre, N. From Integrative Taxonomy to Species Description: One Step Beyond. Syst. Biol. 64, 152–160 (2015).

7. Vakulenko-Lagun, B., Mandel, M. & Betensky, R. A. coxrt: Cox Proportional Hazards Regression for Right-Truncated Data. v. 1.0.2 (2019).

8. Mantel, N. & Myers, M. Problems of convergence of maximum likelihood iterative procedures in multiparameter situations. J. Am. Stat. Assoc. (1971). doi:10.1080/01621459.1971.10482289

9. Collen, B., Purvis, A. & Gittleman, J. L. Biological correlates of description date in carnivores and primates. Glob. Ecol. Biogeogr. 13, 459–467 (2004).

10. Diniz-Filho, J. A. F. et al. Macroecological correlates and spatial patterns of anuran description dates in the Brazilian Cerrado. Glob. Ecol. Biogeogr. 14, 469–477 (2005).

11. Moura, M. R. et al. Geographical and socioeconomic determinants of species discovery trends in a biodiversity hotspot. Biol. Conserv. 220, 237–244 (2018).

12. Gaston, K. J., Blackburn, T. M. & Loder, N. Which species are described first? The case of North-American butterflies. Biodivers. Conserv. 4, 119–127 (1995).

13. Colli, G. R. et al. In the depths of obscurity: Knowledge gaps and extinction risk of Brazilian worm lizards (Squamata, Amphisbaenidae). Biol. Conserv. 204, 51–62 (2016).

14. Essl, F., Rabitsch, W., Dullinger, S., Moser, D. & Milasowszky, N. How well do we know species richness in a well-known continent? Temporal patterns of endemic and widespread species descriptions in the European fauna. Glob. Ecol. Biogeogr. 22, 29–39 (2013).

15. Bebber, D. P. et al. Herbaria are a major frontier for species discovery. Proc. Natl. Acad. Sci. 107, 22169–22171 (2010).

16. Raposo, M. A. et al. What really hampers taxonomy and conservation? A riposte to Garnett and Christidis (2017). Zootaxa 4317, 179 (2017).

17. Dayrat, B. Towards integrative taxonomy. Biol. J. Linn. Soc. 85, 407–415 (2005).

18. Isaac, N. J. B., Mallet, J. & Mace, G. M. Taxonomic inflation: its influence on macroecology and conservation. Trends Ecol. Evol. 19, 464–469 (2004).

19. Garnett, S. T. & Christidis, L. Taxonomy anarchy hampers conservation. Nature 546, 25–27 (2017).

20. Bremer, K., Bremer, B., Karis, P. & Källersjö, M. Time for change in taxonomy. Nature 343, 202–202 (1990).

21. Köhler, J. et al. New Amphibians and Global Conservation: A Boost in Species Discoveries in a Highly Endangered Vertebrate Group. Bioscience 55, 693 (2005).

22. Guedes, J. J. M., Feio, R. N., Meiri, S. & Moura, M. R. Identifying factors that boost species discoveries of global reptiles. Zool. J. Linn. Soc. in press (2020). doi:10.1093/zoolinnean/zlaa029

23. Burgin, C. J., Colella, J. P., Kahn, P. L. & Upham, N. S. How many species of mammals are there? J. Mammal. 99, 1–14 (2018).

24. Mallet, J. Species, Concepts of. in *Encyclopedia of Biodiversity* 6, 679–691 (Elsevier, 2013).

25. Kutner, M. H., Nachtsheim, C. J., Neter, J. & Li, W. Applied Linear Statistical Models. (McGraw-Hill Irwin, 2004).

